# Age-related trajectories of DNA methylation network markers: a parenclitic network approach to a family-based cohort of patients with Down Syndrome

**DOI:** 10.1101/2020.03.10.986505

**Authors:** M. Krivonosov, T. Nazarenko, M.G. Bacalini, M.V. Vedunova, C. Franceschi, A. Zaikin, M. Ivanchenko

**Affiliations:** Research Center for Trusted Artificial Intelligence, Institute for System Programming of the Russian Academy of Sciences, Moscow, 109004, Russia; Department of Applied Mathematics and Laboratory of Systems Medicine of Aging, Lobachevsky University, Nizhny Novgorod, 603950, Russia; Department of Mathematics and Institute for Women’s Health, University College London, London, WC1H 0AY, UK; Department of Experimental, Diagnostic and Specialty Medicine, University of Bologna, Bologna, 40126, Italy; Center for Analysis of Complex Systems, Sechenov First Moscow State Medical University, 119146, Moscow, Russia

**Author notes:** Equally contributed, joint first author (M. Krivonosov, T. Nazarenko).

**Keywords:** Down syndrome, parenclitic network, DNA methylation, complex networks, aging

## Abstract

Despite the fact that the cause of Down Syndrome (DS) is well established, the underlying molecular mechanisms that contribute to the syndrome and the phenotype of accelerated aging remain largely unknown. DNA methylation profiles are largely altered in DS, but it remains unclear how different methylation regions and probes are structured into a network of interactions. We develop and generalize the Parenclitic Networks approach that enables finding correlations between distant CpG probes (which are not pronounced as stand-alone biomarkers) and quantifies hidden network changes in DNA methylation. DS and a family-based cohort (including healthy siblings and mothers of persons with DS) are used as a case study. Following this approach, we constructed parenclitic networks and obtained different signatures that indicate (i) differences between individuals with DS and healthy individuals; (ii) differences between young and old healthy individuals; (iii) differences between DS individuals and their age-matched siblings, and (iv) difference between DS and the adult population (their mothers). The Gene Ontology analysis showed that the CpG network approach is more powerful than the single CpG approach in identifying biological processes related to DS phenotype. This includes the processes occurring in the central nervous system, skeletal muscles, disorders in carbohydrate metabolism, cardiopathology, and oncogenes. Our open-source software implementation is accessible to all researchers. The software includes a complete workflow, which can be used to construct Parenclitic Networks with any machine learning algorithm as a kernel to build edges. We anticipate a broad applicability of the approach to other diseases.

## Introduction

Epigenetic modifications are chemical and/or physical changes in chromatin that can be preserved across cell divisions, and DNA methylation is the most studied among them [1]. This is in part due to the fact that this molecular layer tends to be relatively stable and can be assessed by cost-effective approaches, like the Infinium microarrays, that allow measuring DNA methylation of hundreds of thousands of CpG loci with single base resolution at the genome-wide level [2]. In addition, a bulk of literature demonstrated that DNA methylation changes occur both in physiological (development, sex differentiation, or aging) and pathological (cancer, age-related diseases, or genetic syndromes) conditions [1,3,4].

So far, most of the studies that investigate DNA methylation changes associated with a certain condition focus on individual CpG sites (differentially methylated positions) or groups of chromosomally adjacent CpG sites (differential methylated regions) [5,6]. However, the epigenetic regulation of the genome is a complex and highly integrated system, in which DNA methylation modules are not isolated [7]. Analytical approaches that consider the cross-talk between DNA methylation alterations in different regions of the genome could therefore provide new insights into the epigenetic remodeling occurring in various physiological and pathological conditions, as it has been recently demonstrated [8–11].

In this light, the methods based on network analysis have the potential to provide deeper insights into the regulation of DNA methylation profiles. Among others, the parenclitic network approach [12] is of particular interest, as it enables researchers to represent methylation data in the form of a network, even if functional links between different methylation probes or regions are not established.

Briefly, this approach implements networks, the nodes in which correspond to the relevant parameters, and assigns edges between nodes, if their values significantly deviate from the appropriately defined Control set (group). A measure of deviation is usually introduced on a plane of two coordinates that correspond to the pair of nodes, e.g. Zanin et al. used the distance between a data point to the linear regression line built on a Control group [12]. Since that pioneering work was published, parenclitic networks have been successfully been applied to detecting key genes and metabolites in the context of different diseases, see [13] for a review and [14] for a discussion of applications in brain research. In [15], we have applied this methodology for machine learning classification of human DNA methylation data carrying signatures of cancer development. Later [16], acknowledging that the interactions of two features (at least in biological systems of biomarkers) often cannot be described by a linear model, it was proposed to use 2-dimensional kernel density estimation (2DKDE) to model the Control distribution.

The main difficulty in using such approaches is implementing an end-to-end analysis package (incorporating many technically-complex nested steps), in particular, avoiding *ad hoc* choices of deviation measures and subjective definition of “cut-off thresholds” to discriminate small and large deviations in edge assignment. Here, we present a solution to these issues that employs machine learning algorithms for parenclitic network construction.

We demonstrate the validity of the method for analyzing DNA methylation changes in patients with Down Syndrome (DS). Despite the fact that the cause of DS is well established, the underlying molecular mechanisms of associated epigenetic modifications remain poorly understood. Previous studies have shown that Down Syndrome (DS), which is caused by a full or partial trisomy of chromosome 21, is characterized by a profound remodeling of DNA methylation patterns, which involves not only chromosome 21 but is widespread across the genome [17–20]. DS is a segmental progeroid syndrome, as it is characterized by a phenotype of premature/accelerated aging that occurs in a subset of organs and systems, which include the immune and the nervous systems [21–22]. Based on an epigenetic clock of 353 CpG sites [9], Horvath et al demonstrated accelerated epigenetic aging in DS against age-matched Controls manifested in blood and brain tissues [23].

Among the different cohorts investigated so far, the GSE52588 dataset is of particular interest, as it is a family-based cohort that allows evaluating not only the epigenetic remodeling in DS, but also the potential contribution of genetic background and environment, which are at least in part shared by members of a family [17]. The analysis that focused on differential methylation probes and regions on this dataset highlighted epigenetic alterations in genes involved in the developmental functions (including hematological and neuronal development), metabolic functions, and regulation of chromatin structure, which are in line with the results of independent cohorts [17–20].

In this paper, we aim to go beyond individual or local region differential CpG methylation analysis for Down Syndrome patients and their family members and obtain further insights into the co-differential methylation of CpG pairs and parenclictic network analysis. In particular, we address complex topological changes of the epigenome in disease. In other words, these are the changes, which can be very different for particular DNA methylation probes for individuals, but share a similar topological profile distinct from that of healthy subjects. We also identify genes and molecular pathways associated with the differential network DNA methylation in Down Syndrome not found during individual or local region CpG analysis.

Secondly, (this information is mainly presented in Supporting Information) we present the implementation of an end-to-end workflow to construct Parenclitic Networks, which is available for download; we demonstrate that machine learning algorithms can be chosen as a kernel to build edges in the network using geometric and probabilistic approaches as examples. We also present a new approach (PDF-adaptive) that allows for an automated choice of the “cut-off threshold” to remove insignificant edges.

The paper is structured as follows. First, we apply our algorithm to investigate the age-dependent network signatures of Down Syndrome on methylation data. In particular, we report the signature of the pure DS disease (regardless of age); the signature of age-related changes in a healthy population (which indicates the proximity of DS patient cohort to an older healthy population); the signature, which changes with age in a healthy population slower than in DS patients; and a signature, which changes in a healthy population with age faster than in DS patients. We also conducted the Gene Ontology and KEGG analysis and found associations with the nervous system, cell fate and oncogenesis, cellular communication, development of metabolic syndrome, and female sex hormones.

## Materials and Methods

### Down Syndrome methylation data

We consider a publicly available dataset (GSE52588) as an application and validation of our approach, where the whole blood DNA methylation was assessed by the Infinium HumanMethylation450 BeadChip in a cohort including individuals affected by Down Syndrome (DS), their unaffected siblings (DSS), and their mothers (DSM) [20] (29 families in total).

Age distribution of subjects from the 3 groups is shown in Fig.1(B). As depicted in Fig.1(A), this family-based model (which minimizes potential confounding effects, as members of the same family tend to share the same habits/environment and genetic background) allows for four (4) different comparisons, in which 2 phenotypes (DS and aging) are combined in different ways:

1. comparing DS against both DSM and DSS yields a healthy phenotype;
2. comparing DSS and DSM provides insights into the epigenetic remodeling occurring during aging in euploid subjects;
3. comparing DS and DSS provides insights into the epigenetic remodeling associated with the syndrome in the age-matched groups;
4. comparing DS and DSM discriminates the effects of the syndrome and aging.

**Figure 1.**
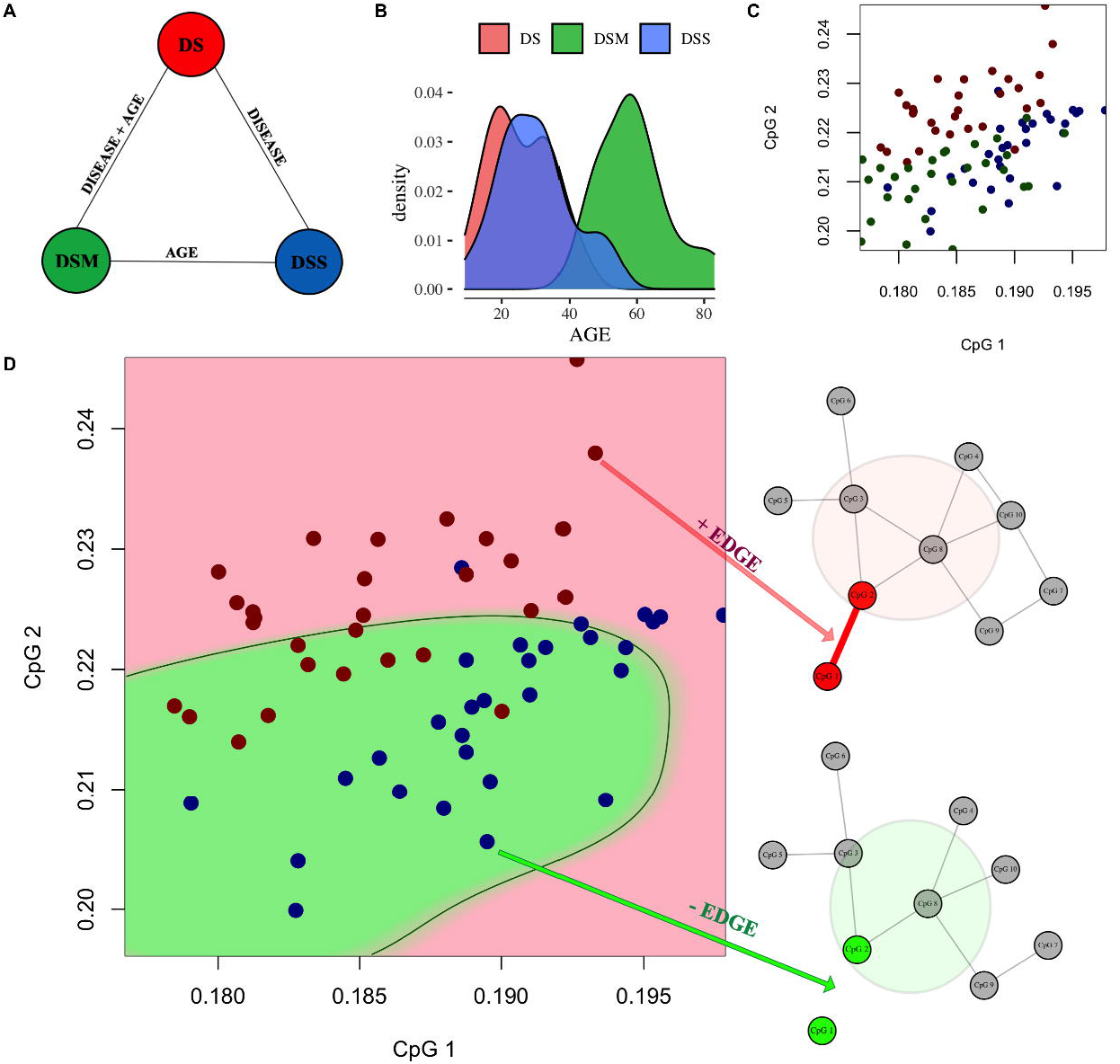
Features of the initial data and the scheme of networks construction. **(A)** Schematic representation of the three groups in the GSE52588 dataset and their phenotypic difference; **(B)** Age-distribution for DSM/DSS/DS groups; **(C)** Example of the three groups representation on the plane of a pair of CpGs loci; **(D)** Illustration of the parenclitic network construction: for instance, on (C) we considered an example of M-control-network, where DSM (green points) is the *Control* group and DS (red points) is the *Case* group and DSS (blue points) is the *Test* group. Using PDF-adaptive (the best threshold) method we detect an area (green Fig. on D), which best describes the area of *Controls*. If the separation accuracy between the *Control* and *Case* groups is less than 90%, the edge between the network nodes (*CpG1, CpG2*) is not assigned for all subjects. Otherwise, the edge (*CpG1, CpG2*) is assigned for a particular subject if it is classified as the *Case* group (if the point falls into the red area). Finally, having repeated the procedure for all pairs of CpG probes, we arrive at individual networks for each subject such that nodes correspond to probes and edges between them designate that a subject falls out of the Control group with regard to methylation of a particular pair of CpGs.

### Preprocessing

Raw *.idat* files were extracted and pre-processed using *minfi* package [24]. Data were normalized using the *preprocessFunnorm* function, while the probes with p-value detection above 0.05 in more than 1% of samples were removed. Furthermore, cross-reactive and polymorphic probes, as reviewed by Zhou et al [25], and probes on the X and Y chromosomes were removed. Finally, the procedure left 412993 probes from the original array. As the entire list of probes was too large for our algorithms, we decided to focus on the subset of probes mapped in CpG Islands and Shores, as previous studies suggest that DNA methylation de-regulation is more likely to occur in these regions (CIT The human colon cancer methylome shows similar hypo- and hypermethylation at conserved tissue-specific CpG island shores [26]). This subset includes 114674 probes. Additionally, we exclude sex-specific CpGs to decrease possible bias relying on the list of sex-specific CpGs [27] with 5665 items. Lastly, the list contains 113630 CpGs (full list of such CpGs can be found in SM-files, in *full_CpGs_list.xlsx* file).

### Generalized Parenclitic Network Analysis

The implementation of a complete workflow to construct Parenclitic Networks is available at https://github.com/mike-live/parenclitic (see Supporting information for a detailed description). The analysis is intended to grasp co-occurring differential changes in DNA methylation of the pairs of probes. We begin by opting out the probes that individually are good separators of the Control and Case groups. That is, for each CpG probe we build a split border classifier based on maximizing the information gain (for details see paragraph ‘Feature Selection Algorithm’ in Supporting Information), and move those that provide accuracy of more than 75% to separate lists (they can be found in SM-files on corresponding sheets (*‘AGE-control-network’, ‘DS-control-network’, ‘S-control-network’, ‘M-control-network’*) in *1d_cpgs.xlsx* file). Such probes are further excluded from the set of nodes of parenclitic networks, and the results are further considered through the prism of single CpG and co-occurring differential CpG methylation.

Overall, we consider four model types of Parenclitic networks processing DS, DSS and DSM groups in different Control, Case, and Test combinations (see illustration on Fig.6 in Supporting information for more details). Here, we employ the conventional machine learning approach, where the groups chosen as Control and Case are used for building classification/discrimination rules, which are subsequently applied to the independent Test group.

1. **DS-Control Network —** the Parenclitic network model of healthy phenotypes, where DS group is the *Control* group, DSS and DSM groups are *Case* groups;
2. **AGE-Control Network —** the Parenclitic network model of aging, where the DSS group is the *Control* group, the DSM group is the *Case* group, the DS group is the *Test* group;
3. **S-Control Network —** the Parenclitic network model of the syndrome, where the DSS group is the *Control* group, the DS group is the *Case* group, the DSM group is the *Test* group;
4. **M-Control Network —** the Parenclitic network model of aging and syndrome, where the DSM group is the *Control* group, the DS group is the *Case* group, the DSS group is the *Test* group.

Parenclitic networks for individual subjects are constructed according to the following rules. For each pair of CpG sites (*CpGj CpGj*), we place subjects on a corresponding two-dimensional plane according to beta-values and mark *Control* and *Case* groups. Fig.1(C,D) illustrates the parenclitic network construction: in (C), we considered example of M-control-network, where DSM (green points) is the *Control* group and DS (red points) is the *Case* group and DSS (blue points) is the *Test* group. Using PDF-adaptive (the best threshold) method, we detect an area (green area Fig.1(D)), which best describes the area of *Controls*. If the separation accuracy between the *Control* and *Case* groups is less than 90%, the edge between the network nodes (*CpGi, CpGj*) is not assigned for all subjects. Otherwise, the edge (*CpGi, CpGj*) is assigned for a particular subject if it is classified as the *Case* group (if the point falls into the red area). Finally, having repeated the procedure for all pairs of CpG probes, we arrive at individual networks for each subject such that nodes corresponding to probes and edges between them designate that a subject falls out of the Control group with regard to methylation of a particular pair of CpGs.

In addition to the obtained networks for each patient separately, we also build “general” networks for each DS/AGE/S/M-Control construction. In each model, we leave only those edges that correspond to separation between Cases and Controls with at least 90% accuracy. In the Fig.2(B), we present resized images of such general DS/AGE/S/M-networks (enlarged images can be accessed via the links given in the caption to the sfigure); constructions are provided in SM-files on corresponding sheets (*‘AGE-control-network’, ‘DS-control-network’, ‘S-control-network’, ‘M-control-network’*) in *union_networks.xlsx*, and lists of vertices of networks can be found in SM-files: in *lists_of_networks_CpGs.xlsx*). The size of each vertex in this representation is associated with the degree (presented in *lists_of_networks_CpGs.xlsx* as a second column) of the vertex in the network (that is, the sum of the edges it contains). Thus, the larger the vertex, the more often this CpG-site has participated in the “successful” separation of Cases and Controls in a pair with another CpG-site. For the top 10 CpGs for each network, we have placed links reporting whether these sites have been previously reported as being related to age [28–30] or related to DS in whole blood [17]. Fig.2(A) shows the Venn Diagram for the vertices (CpG sites) of networks. It demonstrates that each general network is predominantly unique. Further, for each of these generalizing networks (based on the vertices selected in them), we will carry out GO and KEGG analysis and identify biological processes that correspond to differential methylation in Cases and Controls.

**Figure 2.**
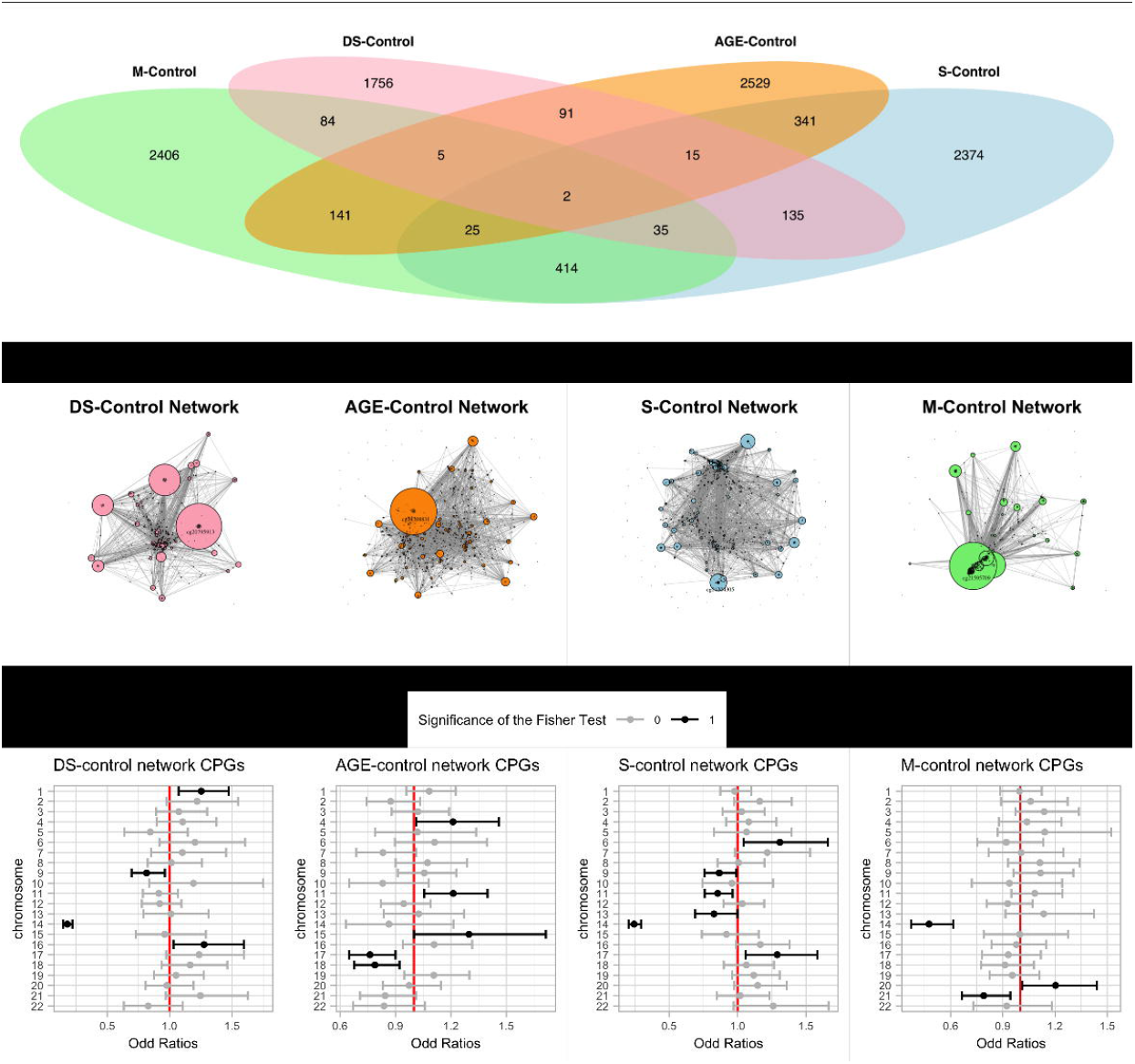
Commonality and individuality of the networks of the four considered models. **(A)** Venn Diagram for the vertices (CpG sites) of generalizing networks; size of nodes in each image **(B)** DS-Control generalizing Network, AGE-Control generalizing Network, S-Control generalizing Network and M-Control generalizing Network are associated with their degree (the greater the degree of the vertex, the more connections it has). The most powerful node (with maximum degree in the network was highlighted by label). Enlarged presentations of networks can be found at https://tatiananazarenko.github.io/PN-DS/. **(C)** The prevalence of CpGs from each chromosome in the network set. We demonstrate here that CpGs from chromosome 21 are not dominant in such networks (most of these chromosome 21 CpGs were selected for 1D analysis and were not included in the parenclitic). It seems important for us to note this fact, since parenclitic analysis was able to capture deeper and more complex relationships between the data.

### Cross-Valida tion

Despite the fact that all further analysis will be carried out on training networks using all the samples in Case and Control groups (to find the signature itself and to further study the behavior of the Test group on it), we carried out an additional cross-validation procedure to show the adequacy of the method as a simple classifier. We did not find other datasets of a similar structure (for example, the well-known GSE107211 dataset studies infants (healthy and DS), and the GSE63347 dataset examines brain tissues). Moreover, the sample size within our dataset is quite small to conduct any standard method of cross-validation (with the division of a group of samples into several folds). However, we resorted to a modification of the Leave-One-Out Cross-Validation (LOOCV) procedure, which, in our case, was called the Leave-(Family)-Out Cross-Validation procedure. Its essence lies in the fact that we repeatedly (29 times, according to the number of families in our set) excluded members of one family from the dataset (one mother and two of her children: a healthy child and a child with DM); trained the networks on the remaining samples and then obtained a prediction (networks) for the members of the discarded family. After such a procedure was carried out for each family, we collected the resulting networks into one set (in each network design), counted the characteristics and calculated the area under the ROC curve (AUC) to see how well the Controls and Cases were divided by such characteristics. We note that in the vector of such predictions (characteristics of networks), the prediction for each sample was obtained on those models, in which the sample itself did not participate in the training process. Fig. 4 (A) shows AUCs for each characteristic for each network design. We note here that a lot of characteristics of DS-Control generalizing Network, S-Control generalizing Network, and M-Control generalizing Network showed excellent performance. The performance of the characteristics of the AGE-Control generalizing Network was noticeably lower, but from our point of view, it was due to the fact that in this model two classes (Control and Cases) were DSS group and DSM group, and their samples slightly overlapped by age (see green and blue distributions in Fig. 1 (B)). From our point of view, this result only additionally confirms the adequacy of the applied approach. Some characteristics show a high quality of class separation, and some - very low. For example, low performance for the characteristic *number of vertices* is easily explained: since in each network design all samples always have the same number of nodes (some of which are connected by edges, and some are not), this characteristic is always equal to the same number and cannot have some class separability property. To highlight the best characteristics across all networks (in terms of their performance), we visualize the left panel with boxplots (where each boxplot stands for AUCs values of networks) and separate those of them, the median value of which exceeds 0.9. Finally, we sort the characteristics by the median of their AUC and highlight those that we will use in further analysis (Fig. 4 (B)).

**Figure 3.**
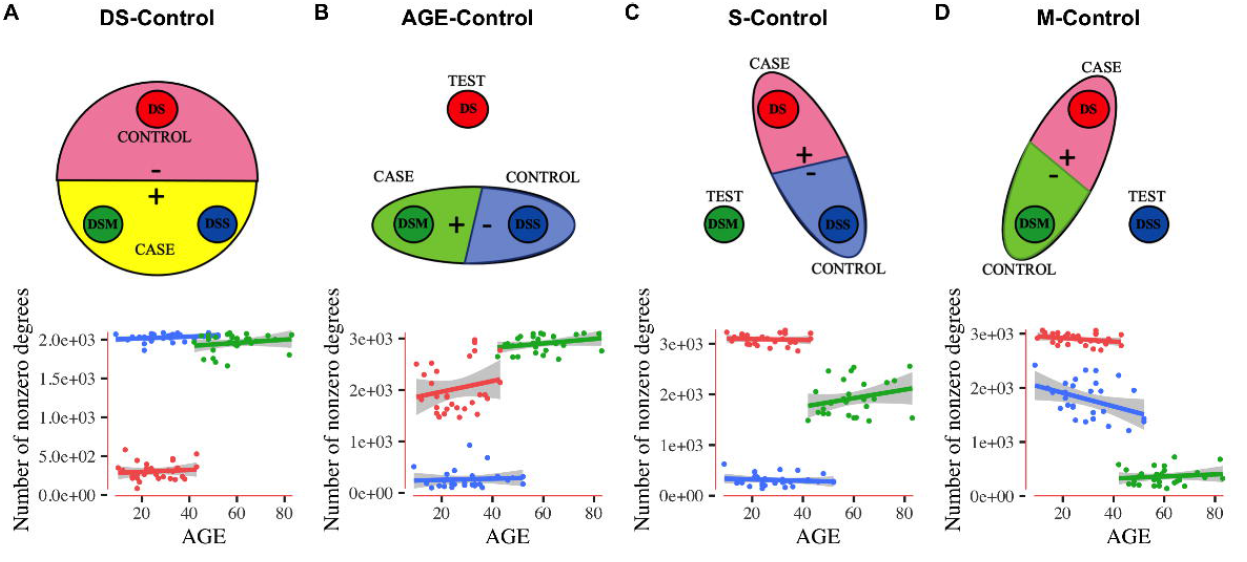
Age dependence of network characteristics. Top panel: Assigning the cohorts for network construction. Bottom panel: Considering the Test group against the Controls and Cases reveals age-related trajectories quantified by network signatures (B-D). Here, we exemplify it in the number of non-zero-degree nodes (the number of CpG probes, that show differential network methylation) in individual networks for DS/AGE/S/M-network design versus AGE (A–D labels respectively). Commonly, the Case group networks are characterized by a considerable number of co-occurring differential CpG methylation changes, as compared to the Control group (as follows from the network construction algorithm). Specifically, for different Control, Case and Test groups: B) Parenclitic network model of aging (AGE-Control): DS samples are closer to DSM than to their DSS and get even closer with age; C) Parenclitic network model of the syndrome: DSM samples get closer with age to DS; D) Parenclitic network model of aging and syndrome: DSS samples get closer with age to DSM. Cf. Supporting Information for the plots for the other network indexes that confirm the described effects.

**Figure 4.**
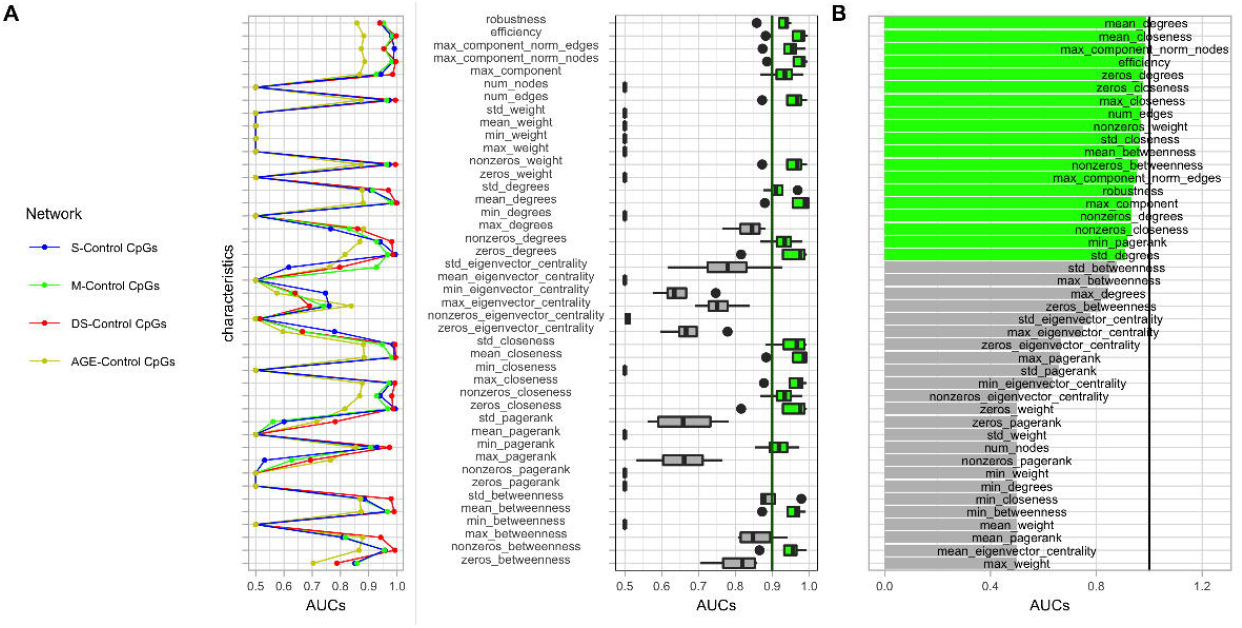
Results of L-(Family)-OUT procedure (A) Left panel: AUCs on each characteristic for each Network design are demonstrated. We note here that a lot of characteristics of DS-Control generalizing Network, S-Control generalizing Network, and M-Control generalizing Network showed excellent performance. The performance of the characteristics of the AGE-Control generalizing Network is noticeably lower, but from our point of view, it is due to the fact that in this model two classes were DSS group and DSM group, and their samples slightly overlapped by age (see green and blue distributions in Fig. 1 (B)). We believe that this result only additionally shows the adequacy of the applied approach. Some characteristics show a high quality of class separation, and some – very low. For example, low performance for the characteristic number of vertices is easily explained: since in each network design all samples always have the same number of nodes (some of which are connected by edges, and some are not), this characteristic is always equal to the same number and cannot have some class separability property. **(A) Right panel:** To highlight the best characteristics across all networks (in terms of their performance), we visualize the left panel with boxplots (where each boxplot is for AUCs values on for networks) and separate those of them whose median value exceeds 0.9. **(B)** Finally, we sort the characteristics by the median of their AUC and highlight those that we will use in further analysis.

We note that the main goal of this work was not to build a classifier but to find the CpGs signature in each network design and further study it using network characteristics of the Test group. However, we consider it important to provide the results of this cross-validation to prove the adequacy of the application of our method and demonstrate that it is not prone to overfitting.

### DNAmAGE and epigenetic age acceleration

DNAmAGE (DNA methylation age) was calculated using the online tool available at https://dnamage.genetics.ucla.edu/. Epigenetic age acceleration values (residuals) were calculated as the difference between the Horvath’s epigenetic age and chronological age.

### Functional enrichment analysis

Gene Ontology enrichment was performed using the *methylgometh* function implemented in the methylGSA Bioconductor package [31]. Two kinds of input data were used: the lists of single CpGs differentially methylated between the groups of Cases and Controls (one-dimensional separators) and the lists of CpGs that constitute parenclitic networks (parenclictic network separators). The results are presented in Table 2 (for Parenclitic) and Table 3 (for 1D), Supporting Information. As the parenclitic approach does not return an explicit p-value, the input file for *methylgometh* function was created by assigning value less than its cut-off to all the CpG sites selected by the parenclitic analysis, and value greater than cut-off to the other CpG sites. REVIGO (http://revigo.irb.hr/) was used to remove redundant GO terms. Functional annotation analysis was performed using the ‘Functional Annotation Charts’ tools of the Database for Annotation, Visualization and Integrated Discovery (DAVID Bioinformatics Resources 6.8, NIAID/NIH) [d_7]. The enrichment analysis involved the lists of genes that contain differentially methylated CpGs revealed by the parenclitic approach. The identifier “official gene symbol” (Homo sapiens) was used as an annotation category, and KEGG_PATHWAYS was used for the pathway analysis. Significantly enriched pathways (from the KEGG_PATHWAYS database) with Benjamini < 0.05 and P < 0.05 were selected. Results of KEGG_PATHWAYS analysis are presented in Table 1, Supporting Information.

## Results

### Network analysis and identification of network-age-dependent effects

We previously demonstrated [20,26] that individuals with DS tend to have a higher epigenetic age compared to age-matched Controls by Horvath’s clocks, and found some signatures of DS based on the local differential DNA methylation analysis. In this work, for the first time ever, we conduct a network analysis of differential methylation in DS taking into account genome-wide associations and provide network DNA methylation signatures of DS based on co-occurrence of probe methylation modifications.

The proposed method allows obtaining individual parenclitic networks for each subject. This way, we analyzed the statistical properties of such networks (betweenness, pagerank, closeness, eigenvector centrality, nodes degrees, number of edges and nodes, maximum connected component, efficiency, robustness) and demonstrated that they differ between the classes (see, for example, Figures 7, 9, 11 and 13 in Supporting Information).

This method also determines signatures (specific CpG sites) associated with each of DS/AGE/S/M-Control models (see Fig.2(B)). These signatures are essentially unique (see Fig.2(A)) for each model. At the same time, the specificity is also observed in different aging trends (for example, in the characteristics of the M-control network, the DS group is closer to young representatives of healthy population, and in the characteristics of the S-network - on the contrary, closer to the elderly) (see Fig.3).

We illustrate findings focusing on the number of non-zero-degree edges in the parenclitic network. This quantity characterizes the degree of difference of DNA methylation patterns in the individual from the Control group in terms of differential network CpG methylation. The observed trends in each model case (especially, in the behavior of the *Test* group), allows the following generalizations (Fig.3) are as follows:

- Healthy phenotype model network (DS-control): here, DSM and DSS show similar networking properties and are very different from DS (see, for example, number of non-zero-degree nodes in Fig.3(A) and other characteristics in Fig.7 in Supporting Information). We assume that the signature of this network (the involved nodes) is responsible for the pure (regardless of age) difference between the healthy population and patients with DS;
- Parenclitic model of aging (Age-control): DS network signatures are closer to DSM than to DSS and get closer with age (see. Fig.3(B) and Fig.9 in Supporting Information). This fact corroborates with the established concept of accelerated aging in DS. Examining the behavior of network characteristics against age acceleration (see, for example number of non-zero-degree nodes in Fig.5 and other characteristics in Fig.9 in Supporting Information), we observe that the greater the absolute age acceleration, the closer patients with DS are to their mothers. Notably, this trend is seen for both the positive and negative age accelerations. The latter could indicate the limitations of Horvath’s epigenetic clocks for some special phenotypes of DS patients, where formally negative age acceleration does not reflect actual accelerated biological aging. In this way, the parenclitic network signatures behave more robustly, consistently indicating the difference between the healthy elderly and the healthy young populations, and the proximity of DS patients to the mothers, especially in the elderly group (indicating a strong effect of age-accelerating for DS patients);
- Parenclitic model of DS (S-control): DSM network signatures are different from DSS, and get closer to DS with age (see, Fig.3(C), and other characteristics in Fig.11 in Supporting Information), which is also in line with the concept of age-acceleration in DS. We infer that the signature of this network (the involved nodes) captures the difference between the DS patients and their siblings. Moreover, the trend in the group of mothers suggests that in a healthy population this signature changes with age slower (and only with increasing age does it approach the state of the DS group);
- Parenclitic model of both aging and syndrome (M-control): DSM network signatures are different from DS and DSS groups. This trend shows that this group moves away from DS peers and gets closer to mothers with age (see, for example the number of non-zero-degree nodes, Fig. 3(D), and other characteristics, Fig. 13 in Supporting Information). We assume that the signature of this network (the nodes selected into it) changes with age faster in the healthy population than in DS patients.

**Figure 5.**
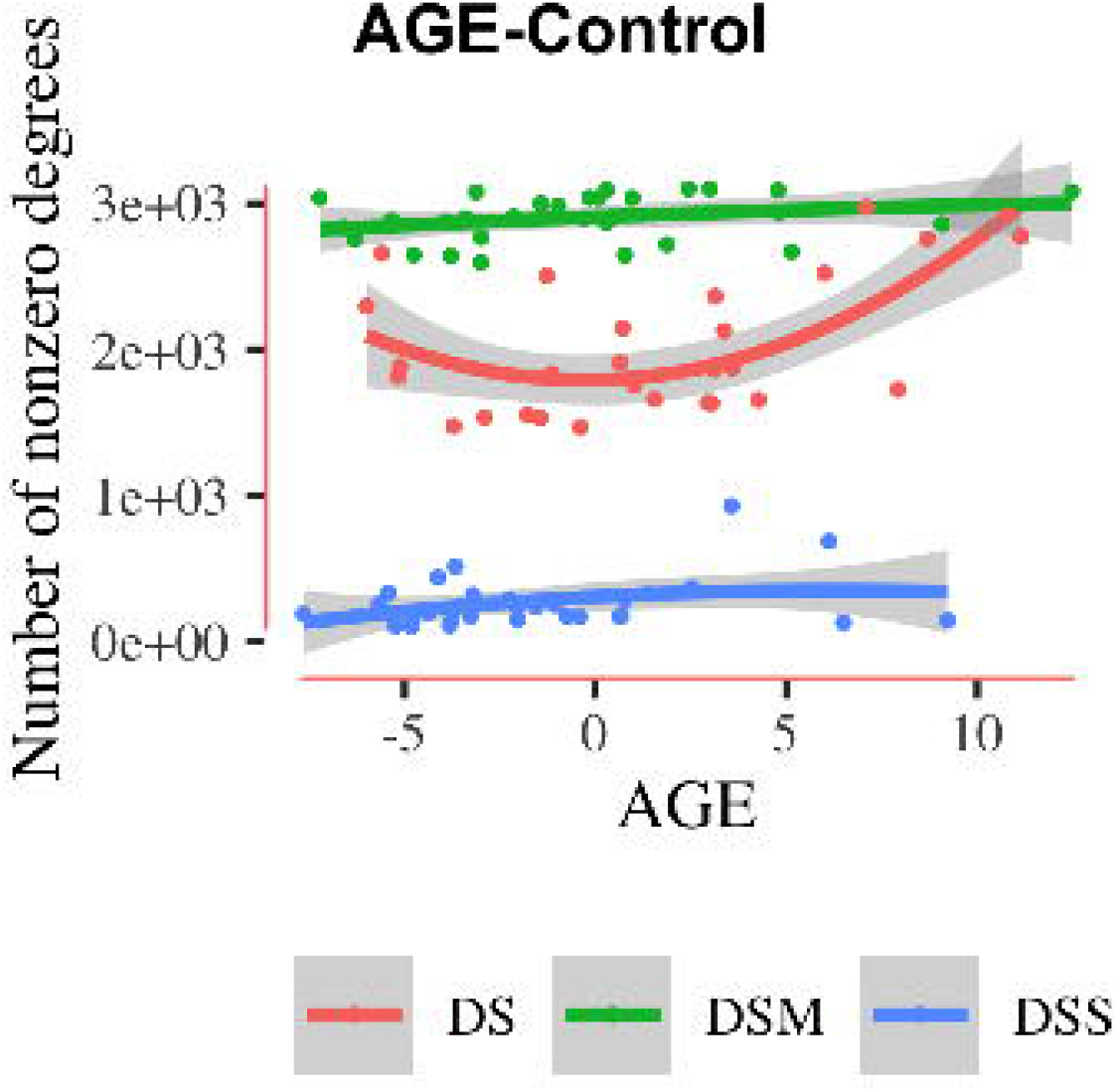
Age-acceleration dependence of network characteristics. The number of nonzero degree network nodes (the number of CpG probes that show differential network methylation) vs age acceleration (horizontal axis). In all cases, the DS methylation network signature is closer to mothers. For the other network characteristics see Supporting Information.

### Gene Ontology and KEGG Results

In this section, we identify genes responsible for age related trajectories of methylation network signatures and make an enrichment analysis in Gene Ontology (GO) terms. Note that we conduct a separate analysis of the lists of CpGs selected by the parenclitical approach and the lists of CpGs identified as good individual (1D) classifiers. The results are summarized in Table 2 (for parenclitic approach) and Table 3 (for 1D approach) in Supporting Information. While the DS-Nodes comparison did not return any significantly enriched GO, 23, 37, and 77 GO terms were enriched in the lists of CpG probes resulting from the S-Nodes comparison, M-Nodes comparison, and AGE-Nodes comparison, respectively. A substantial overlap was observed between these 3 cases with 19 terms in common. To gain more insights into the shared and specific gene ontology, we applied REVIGO to the lists of GO terms to remove redundant terms and focus on the “Biological Processes” description (Table 2, Supporting Information). We found 5 GO terms (pattern specification process, regionalization, cell fate commitment, skeletal system development, appendage development) common to all the comparisons, and another 5 GO terms (embryonic organ morphogenesis, epithelial cell differentiation, forebrain development, camera-type eye development, neuron migration) shared between the S-Nodes comparison and the M-Nodes comparison. Notably, the same GO analysis on the results of 1D comparison (Table 3, Supporting Information) returned a substantially lower number of enriched GOs in the S-Nodes comparison and in the M-Nodes comparison, and no overlap between these two comparisons and the AGE-Nodes comparison.

The earlier study, which was based on the local probe differential DNA methylation analysis and employed the same data set [17], identified some specific methylation changes in individuals with Down syndrome, as compared to their healthy siblings and mothers. These differences mainly concerned the genes responsible for morphogenesis and intracellular processes. Notably, the use of parenclitic analysis made it possible to identify more complex patterns of epigenetic changes occurring in Down syndrome. Analysis in GO revealed significant differences in the methylation of loci associated with the functioning of the nervous system, as well as a large number of processes (8 out of 36) related to the development of the genitourinary system and the musculoskeletal system. Biological processes common to all groups are associated with cell differentiation (GO:0007389 pattern specification process, GO:0003002 regionalization, GO:0045165 cell fate commitment) and morphogenesis (GO:0001501 skeletal system development, GO:0048736 appendage development).

Identified signatures highlight the molecular mechanisms of changes associated with the age development of Down Syndrome. Analysis of the Kyoto Encyclopedia of Genes and Genomes (KEGG) (see (Table 1, Supporting Information)) shows a large number of signaling pathways associated with i) the nervous system (Neuroactive ligand-receptor interaction, Circadian entrainment, Axon guidance, Glutamatergic synapse, Calcium signaling pathway, GABAergic synapse, Morphine addiction, Nicotine addiction, Retrograde endocannabinoid signaling, Oxytocin signaling pathway, Cocaine addiction, Dopaminergic synapse, Glycosaminoglycan biosynthesis - heparan sulfate / heparin, Cholinergic synapse); ii) Cell fate and oncogenesis (Wnt signaling pathway, Signaling pathways regulating pluripotency of stem cells, Pathways in cancer, cAMP signaling pathway, Basal cell carcinoma, Proteoglycans in cancer); iii) Cellular communication (Rap1 signaling pathway, Calcium signaling pathway, Gap junction, PI3K-Akt signaling pathway, Glycosaminoglycan biosynthesis - heparan sulfate / heparin); iv) Development of metabolic syndrome (Adrenergic signaling in cardiomyocytes, Type II diabetes mellitus, Dilated cardiomyopathy, Insulin secretion, Maturity onset diabetes of the young, Arrhythmogenic right ventricular cardiomyopathy (ARVC)); and v) Female sex hormones (Oxytocin signaling pathway, Estrogen signaling pathway). These groups of genes were not identified via local differential DNA methylation analysis [20].

## Discussion

While plenty of results demonstrate and build upon single CpG sites or local regions of differential DNA methylation in the context of pathology or aging or age-related diseases, little is known about the non-local group CpG differential methylation and the corresponding functional pathway enrichment. This study aims to fill this gap by advancing and employing the co-differential methylation analysis for genome-wide pairs of CpGs for Down Syndrome patients and their family members, and, subsequently, to advance and analyze the so-called parenclictic networks of such pairs as nodes connected by edges.

We demonstrate how parenclitic networks can be used to find aging trajectories of methylation network markers in Down Syndrome. Three cohorts of the analyzed data set - Down Syndrome individuals, their Mothers, and Siblings - allowed us to construct different Parenclitic Networks depending on the choice of Control, Case, and Test groups. The pipeline starts with identifying CpG sites, which are good individual classifiers, and pairs of CpGs sites, which are good two-dimensional classifiers, but fail when studied separately. The latter set is used to build a parenclitic network with edges corresponding to significant differential paired CpG methylation. In terms of the structure, the Case networks are much richer in non-zero-degree nodes and edges than Control networks. Test networks do not follow such preset, and their closeness to other groups needs to be determined by network properties and age-related changes. On top of that, we build “general” networks that include only nodes and edges common to all individuals from a group, thus defining an epigenetic network signature and an entry set for further GO analysis.

DS has many well-described clinical manifestations including mental retardation, congenital heart defects, gastrointestinal and hematological abnormalities, and other specific signs that ultimately form complex unique phenotypes. Here, we decomposed the epigenetic signature of DS patients into categories based on the hidden links between the covariates: the signature of the pure DS disease (regardless of age); the signature of age-related changes in a healthy population (which defines a plurality of DS patients as being close to an older population); the signature, which changes with age in a healthy population slower than in DS patients; and the signature, which changes in a healthy population with age faster than in DS patients. This could help, in the future, to identify molecular targets for medical treatment of diseases accompanying DS.

Our study identified network molecular changes associated with the DS development. The molecular changes emerging in the DS support the fact that genes do not act as autonomous units of the genome, but operates as parts of spatially coordinated regulatory networks. Accordingly, epigenetic remodeling does not affect only chromosome 21, but the entire genome. We showed that patients with DS have different epigenetic changes in CpG sites of genes related to the clinical manifestations of DS. These are primarily biological processes responsible for the functioning of the central nervous system, skeletal muscles, disorders in carbohydrate metabolism, cardiopathology, and oncogenes. Instructively, most of the identified epigenetic changes in DS are involved in the same processes as biological pathways undergoing normal age-dependent epigenetic changes. Unique biological processes not associated with age are related to the brain development and organogenesis. DS is characterized by marked changes in brain formation during embryogenesis and in the early postnatal period (reduced cell proliferation, disorganized patterns of cortical stratification, pronounced neurodegeneration after birth, and reduced number of dendritic spines). Most likely, the same molecular mechanisms continue to play a key role in the formation of the phenotype of the central nervous system in adulthood (ex. diminished brain size, smoothed gyrus).

We believe that the networks constructed in this work are not the end of the study, but only the beginning, as the new information obtained in the course of network analysis can be now widely analyzed by biologists and clinicians in order to identify molecular mechanisms resulting in and accompanying DS. As noted in the Cross-Validation section, we could not identify other datasets that could directly validate our results (for example, the well-known GSE107211 dataset studies infants (healthy and with Down syndrome), and the GSE63347 dataset examines brain tissue). Nevertheless, we believe that a separate study of such datasets (finding methylation signatures based on the groups represented in it by a parenclitic approach) and then the correlation of the obtained results with the current ones can be an extremely interesting continuation of our work.

Additionally, this network analysis may help to identify new molecular targets for treatment of patients with DS to prevent their accelerated aging. The networks built in this study require a further detailed analysis and can help researchers involved in DS research to discover new interpretations based on the interactions detected. We particularly highlight the fact that the main idea of the Parenclitic Network approach, the separation of Case-Control group states, can be also applied for analyzing the transition in time, by age or other continuous scale, from one state to another. Introducing the Test set, which is located between the Case and Control states, can become an additional methodological improvement in the parenclitic study of complex systems.

We presented our algorithm as an open-source implementation of Generalized Parenclitic Network analysis to make it more accessible to all researchers. One of the main methodological advances is introducing new machine learning methods and kernels and discussing the possibility of their selection based on the particular features of the problem. We believe that a simple integration into the overall implementation will allow researchers to use not only the proposed methods but also join forces and test their own ideas. We note that we settled on the use of PDF-adaptive, since our data (see the third column in Fig. 6 in the Supporting Information) has a non-linear structure and is better described by the ‘clouds’ of the PDF-adaptive approach (as compared, for example, to the linear SVM method, which divides the plane into two halves with a straight line). We also believe that since the parenclitic kernel uses a classifier that is always based only on a pair of features, the use of more complicated methods may seem redundant and the best methods are those that more accurately capture the control area (any form of it). Together with the PDF-adaptive approach, one could also use the SVM-radial algorithm, which would capture such clouds in a similar way.

The main limitation of the proposed approach is the significant computational time required to build and assess the quality of classifiers based on methylation of each CpG pair due to the huge number of possible pairs. This can be overcome by exploiting parallel calculations on high-performance computers. The current study was also limited by a relatively small number of participants. However, this was mitigated by the structure of the dataset that included close relatives (mothers and siblings) of Down Syndrome patients that ensured the genetic proximity of the participants, as well as the similarity of living and environmental conditions within each family, decreasing the contribution of population heterogeneity.

This design can be applied to any disease, including, e.g., cancer, in which, in addition to critical states, such as a healthy or diagnosed patient, there is data on intermediate conditions. These conditions can be, for example, the diagnosed patients analyzed at an earlier time, when they could be considered healthy. We believe that the results of such an analysis based on Generalized Parenclitic Networks will not only help in the early diagnosis of the disease, e.g., by identifying critical transition marks and risk assessment, but also would shed light on the process itself, through attributes involved in it.

## Acknowledgments

This work was supported by a grant for research centers in the field of artificial intelligence, provided by the Analytical Center for the Government of the Russian Federation in accordance with the subsidy agreement (agreement identifier 000000D730321P5Q0002) and the agreement with the Ivannikov Institute for System Programming of the Russian Academy of Sciences dated November 2, 2021 No. 70-2021-00142. Authors acknowledge non-financial support by the project of the Ministry of Education and Science of the Russian Federation Agreement No. 075-15-2021-639. AZ, TN acknowledge support by the MRC grant MR/R02524X/1.

## Author contributions statement

AZ, CF, MI designed the study, MK developed software, TN developed methodology of age-related analysis, MK and TN conducted the graph analysis, MK, TN, MGB and MV conducted the data analysis. All authors have written and reviewed the manuscript.

## Additional information

Authors declare no competing interests.

## Notes

https://github.com/mike-live/parenclitic

### Competing Interest Statement

The authors have declared no competing interest.

### Summary of Updates

The manuscript was simplified and cross-validation was performed. Author affiliations updated.

## References

[1] Greenberg MVC., Bourc’his D. The diverse roles of DNA methylation in mammalian development and disease. Nat. Rev. Mol. Cell Biol. 2019; 20, 590–607. https://doi.org/10.1038/s41580-019-0159-6.

[2] Bibikova M, Barnes B, Tsan C, Ho V, Klotzle B, Le JM, Delano D, Zhang L, Schroth G.P., Gunderson KL, Fan J-B, Shen R. High density DNA methylation array with single cpg site resolution. Genomics. 2011; 98 (4), 288–295, DOI: https://doi.org/10.1016/j.ygeno.2011.07.007.

[3] Cerrato F, Sparago A, Ariani F, Brugnoletti F, Calzari L, Coppedè F, De Luca A, Gervasini C, Giardina E, Gurrieri F, Lo Nigro C, Merla G, Miozzo M, Russo S, Sangiorgi E, Sirchia SM, Squeo GM, Tabano S, Tabolacci E, Torrente I, Genuardi M, Neri G, Riccio A. DNA Methylation in the Diagnosis of Monogenic Diseases. Genes. 2020; 11(4):355. https://doi.org/10.3390/genes11040355.

[4] Robertson KD. DNA methylation and human disease. Nat. Rev. Genet. 2005; 6, 597–610. https://doi.org/10.1038/nrg1655.

[5] Li D, Xie Z, Le Pape M, Dye T. An evaluation of statistical methods for DNA methylation microarray data analysis. BMC Bioinforma. 2015; 16, 217. https://doi.org/10.1186/s12859-015-0641-x.

[6] Maksimovic J, Phipson B, Oshlack A. A cross-package Bioconductor workflow for analysing methylation array data. F1000Res. 2016 Jun 8; 5:1281. 10.12688/f1000research.8839.3. PMID: 27347385; PMCID: PMC4916993.

[7] Hartwell LH, Hopfield JJ, Leibler S, Murray AW. From molecular to modular cell biology. Nature. 1999; 402, 47–52. https://doi.org/10.1038/35011540.

[8] Cui Z-J, Zhou X-H, Zhang H-Y. DNA Methylation Module Network-Based Prognosis and Molecular Typing of Cancer. Genes. 2019; 10(8):571. https://doi.org/10.3390/genes10080571.

[9] Horvath S, Zhang Y, Langfelder P, Kahn RS, Boks, MPM, Eijk K, Berg LH, Ophoff RA. Aging effects on DNA methylation modules in human brain and blood tissue. Genome Biol. 2012; 13, R97. https://doi.org/10.1186/gb-2012-13-10-r97.

[10] Lund JB, Li S, Baumbach J, Christensen K, Li W, Mohammadnejad A, Pattie A, Marioni RE, Deary IJ, Tan. Weighted gene co-regulation network analysis of promoter DNA methylation on all-cause mortality in old-aged birth cohorts finds modules of high-risk associated biomarkers. J Gerontol A Biol Sci Med Sci. 2020; 75(12), 2249–2257. https://doi.org/10.1093/gerona/glaa066.

[11] Yuan L, Huang, D-S. A network-guided association mapping approach from DNA methylation to disease. Sci Rep. 2019; 9, 5601. https://doi.org/10.1038/s41598-019-42010-6.

[12] Zanin M, Alcazar J, Carbajosa J, Paez MG, Papo D, Sousa P, Menasalvas E, Boccaletti S. Parenclitic networks: uncovering new functions in biological data. Sci. Reports, 2014; (4), 5112. https://doi.org/10.1038/srep05112

[13] Zanin M, Papo D, Sousa P, Menasalvas E, Nicchi A, Kubik E, Boccaletti S. Combining complex networks and data mining: Why and how. Phys. Reports. 2016; 635, 1–44. https://doi.org/10.1016/j.physrep.2016.04.005

[14] Papo D, Buldú JM, Boccaletti S, Bullmore ET. Complex network theory and the brain. Phil. Trans:R.Soc.B. 2014; 369, 1–7. https://doi.org/10.1098/rstb.2013.0520

[15] Karsakov A, Bartlett T, Ryblov A, Meyerov I, Ivanchenko M, Zaikin A. Parenclitic network analysis of methylation data for cancer identification. PloS ONE. 2017; 12, e0169661. https://doi.org/10.1371/journal.pone.0169661

[16] Whitwell HJ, Blyuss O, Menon U, Timms JF, Zaikin A. Parenclitic networks for predicting ovarian cancer. Oncotarget. 2018; 9, 22717–22726. https://doi.org/10.18632/oncotarget.25216.

[17] Bacalini MG, Gentilini D, Boattini A, Giampieri E, Pirazzini C, Giuliani C, Fontanesi E, Scurti M, Remondini D, Capri M, Cocchi G, Ghezzo A, Del Rio A, Luiselli D, Vitale G, Mari D, Castellani G, Fraga M, Di Blasio AM, Salvioli S, Franceschi C, Garagnani P. Identification of a DNA methylation signature in blood cells from persons with Down Syndrome. Aging (Albany NY). 2015 Feb;7(2):82–96. https://doi.org/10.18632/aging.100715. PMID: 25701644; PMCID: PMC4359691.

[18] Do C, Xing Z, Yu YE, Tycko B. Trans-acting epigenetic effects of chromosomal aneuploidies: lessons from down syndrome and mouse models. Epigenomics. 2017; 9, 189–207. https://doi.org/10.2217/epi-2016-0138.

[19] Henneman P, Bouman A, Mul A, Knegt L, van der Kevie-Kersemaekers A-M, Zwaveling-Soonawala N, Meijers-Heijboer HEJ, Trotsenburg ASP, Manners MM. Widespread domain-like perturbations of DNA methylation in whole blood of Down syndrome neonates. PLoS ONE. 2018; 13(3): e0194938. https://doi.org/10.1371/journal.pone.0194938

[20] Laufer BI, Hwang H, Vogel Ciernia A, Mordaunt CE, LaSalle JM. Whole genome bisulfite sequencing of down syndrome brain reveals regional DNA hypermethylation and novel disorder insights. Epigenetics. 2019, 14, 672–684. https://doi.org/10.1080/15592294.2019.1609867.

[21] Franceschi C, Garagnani P, Gensous N, Bacalini MG, Conte M, Salvioli S. Accelerated bio-cognitive aging in Down syndrome: State of the art and possible deceleration strategies. Aging Cell. 2019 Jun;18(3):e12903.. https://doi.org/10.1111/acel.12903 Epub 2019 Feb 15. PMID: 30768754; PMCID: PMC6516152.

[22] Martin GM. Genetic syndromes in man with potential relevance to the pathobiology of aging. Birth Defects Orig Artic Ser. 1978;14(1):5–39. PMID: 147113.

[23] Horvath S, Garagnani P, Bacalini MG, Pirazzini C, Salvioli S, Gentilini D, Di Blasio AM, Giuliani C, Tung S, Vinters HV, Franceschi C. Accelerated epigenetic aging in Down syndrome. Aging Cell. 2015 Jun;14(3):491–5. https://doi.org/10.1111/acel.12325. Epub 2015 Feb 9. PMID: 25678027; PMCID: PMC4406678.

[24] Aryee MJ, Jaffe AE, Corrada-Bravo H, Ladd-Acosta C, Feinberg AP, Hansen KD, Irizarry RA. Minfi: a flexible and comprehensive Bioconductor package for the analysis of Infinium DNA methylation microarrays. Bioinformatics. 2014 May 15;30(10):1363–9. https://doi.org/10.1093/bioinformatics/btu049. Epub 2014 Jan 28. PMID: 24478339; PMCID: PMC4016708.

[25] Zhou W, Laird PW, Shen H. Comprehensive characterization, annotation and innovative use of Infinium DNA methylation BeadChip probes. Nucleic Acids Res. 2017 Feb 28;45(4):e22. https://doi.org/10.1093/nar/gkw967. PMID: 27924034; PMCID: PMC5389466.

[26] Irizarry R, Ladd-Acosta C, Wen B, Wu Z, Montano C, Onyango P, Cui H, Gabo K, Rongione M, Webster M, Ji H, Potash JB, Sabunciyan S & Feinberg AP. The human colon cancer methylome shows similar hypo-and hypermethylation at conserved tissue-specific CpG island shores. Nat Genet 41, 178–186 (2009). https://doi.org/10.1038/ng.298

[27] Singmann P, Shem-Tov D, Wahl S, Grallert H, Fiorito G, Shin S-Y, Schramm K, Wolf P, Kunze S, Baran Y, Guarrera S, Vineis P, Krogh V, Panico S, Tumino R, Kretschmer A, Gieger C, Peters A, Prokisch H, Relton CL, Matullo G, Illig T, Waldenberger M, Halperin E. Characterization of whole-genome autosomal differences of DNA methylation between men and women. Epigenetics & Chromatin. 2015; 8, 43. https://doi.org/10.1186/s13072-015-0035-3

[28] Fernández AF, Bayón GF, Urdinguio RG, Toraño EG, García MG, Carella A, Petrus-Reurer S, Ferrero C, Martinez-Camblor P, Cubillo I, García-Castro J, Delgado-Calle J, Pérez-Campo FM, Riancho JA, Bueno C, Menéndez P, Mentink A, Mareschi K, Claire F, Fagnani C, Medda E, Toccaceli V, Brescianini S, Moran S, Esteller M, Stolzing A, de Boer J, Nisticò L, Stazi MA, Fraga MF. H3K4me1 marks DNA regions hypomethylated during aging in human stem and differentiated cells. Genome Res. 2015 Jan;25(1):27–40. https://genome.cshlp.org/lookup/doi/10.1101/gr.169011.113. Epub 2014 Sep 30. Erratum in: Genome Res. 2019 Apr;29(4):710.2. PMID: 25271306; PCMID: PMC4317171.

[29] Yusipov I, Bacalini MG, Kalyakulina A, Krivonosov M, Pirazzini C, Gensous N, Ravaioli F, Milazzo M, Giuliani C, Vedunova M, Fiorito G, Gagliardi A, Polidoro S, Garagnani P, Ivanchenko M & Franceschi C. Age-related DNA methylation changes are sex-specific: a comprehensive assessment. Aging (Albany NY). 2020; 12:24057–24080. https://doi.org/10.18632/aging.202251

[30] Steegenga WT, Boekschoten MV, Lute C, Hooiveld GJ, de Groot PJ, Morris TJ, Teschendorff AE, Butcher LM, Beck S, Müller M. Genome-wide age-related changes in DNA methylation and gene expression in human PBMCs. Age (Dordr). 2014 Jun;36(3):9648. https://link.springer.com/article/10.1007/s11357-014-9648-x. Epub 2014 May 2. PMID: 24789080; PCMID: PMC4082572.

[31] Ren X, Kuan PF. methylGSA: a Bioconductor package and shiny app for DNA methylation data length bias adjustment in gene set testing. Bioinformatics. 2019; 35, 1958–1959. https://doi.org/10.1093/bioinformatics/bty892

