## Supporting Information for "Age-related trajectories of DNA methylation network markers: a parenclitic network approach to a family-based cohort of patients with Down Syndrome"

### Supplementary Figure Legends

**Figure 6. (A-(M/S/AGE/DS))** Definition of Control and Case groups; **(B-(M/S/AGE/DS))** Illustration of the construction of the Control-area by Control group in each design and the automatic determination of the best border (accuracy of separation of Control and Case groups); **(C-(M/S/AGE/DS))** Illustration networks for members of one family: DSM (green network), DSS (blue network), DS (red network. Participants of the Control group are characterized by a discharged network structure; participants of the Case group are characterized by a dense and connected network structure, and participants of the Test group (for the first two networks) are characterized by an intermediate state (between discharged and density).

**Figure 7. (A)** Characteristics of individual networks in DS-network design versus AGE. As expected, the healthy (DSS and DSM) and DS groups are clearly distinguishable by most characteristics (since the very construction of networks ensures the receipt of different topological objects for Control and Case groups). **(B)** Characteristics of individual networks in DS-network design versus RESIDUALS.

**Figure 8.** Individual networks in the DS-network design grouped into families and ordered in three different ways (by age of each **(A)** DSM, **(B)** DSS, **(C)** DS group).

**Figure 9. (A)** Characteristics of individual networks in AGE-network design versus AGE. As expected, the DSS and DSM groups are clearly distinguishable by most characteristics (since the very construction of networks ensures the receipt of different topological objects for Control and Case groups). At the same time, the Test group (DS) lies closer to DSM group (healthy population older than them) than to DSS (their healthy peers). **(B)** Characteristics of individual networks in AGE-network design versus RESIDUALS.

**Figure 10.** Individual networks in the AGE-network design grouped into families and ordered in three different ways (by age of each **(A)** DSM, **(B)** DSS, **(C)** DS group)

**Figure 11. (A)** Characteristics of individual networks in S-network design versus AGE. As expected, the DSS and DS groups are clearly distinguishable by most characteristics (since the very construction of networks ensures the receipt of different topological objects for Control and Case groups). At the same time, the Test group (DSM) demonstrates the transition in age from the group of DSS (healthy population younger than them) to the DS group (unhealthy population younger than them).**(B)** Characteristics of individual networks in S-network design versus RESIDUALS

**Figure 12.** Individual networks in the S-network design, grouped into families and ordered in three different ways (by age of each **(A)** DSM, **(B)** DSS, **(C)** DS group)

**Figure 13. (A)** Characteristics of individual networks in M-network design versus AGE. As expected, the DSM and DS groups are clearly distinguishable by most characteristics (since the very construction of networks ensures the receipt of different topological objects for Control and Case groups). At the same time, the Test group (DSS) demonstrates the transition in age from the group of DS (their unhealthy peers) to the group of DSM (healthy population older than them). **(B)** Characteristics of individual networks in M-network design versus RESIDUALS.

**Figure 14.** Individual networks in the M-network design grouped into families and ordered by three different ways (by age of each **(A)** DSM, **(B)** DSS, **(C)** DS group)

**Figure 15.** Average number of identified edges in all networks for DS-Control depending on the *minimal score* (see paragraph *Thresholds analysis* in Supporting Information)

**Figure 16.** Number of nodes with non-zero degrees of individual networks (DS-Control) depends on age for three different parameter values of *minimal score* (0.85, 0.9, 0.95).

**Figure 17.** Average number of identified edges in all networks for S-Control depends on the *1-dimensional maximal score* (MS1D) (see paragraph *Thresholds analysis* in Supporting Information)

**Figure 18.** Number of nodes with non-zero degrees of individual networks (S-Control) depends on age for three different parameter values of *1-dimensional maximal score* (MS1D: 0.7, 0.76, 0.8).

### Supplementary Tables Legends

**Table 1.** Kyoto Encyclopedia of Genes and Genomes (KEGG) results (p value<0.05 and Benjamini <0.05).

**Table 2.** Gene Ontology results for Nodes of S/M/DS/AGE-Control Networks. Several processes that are significant for the S-Control and M-Control Networks at the same time (e.g. *pattern specification process, regionalization, cell fate commitment, skeletal system development, appendage development*) turn out to be significant also for the AGE-Control (which, on a par with the study of network characteristics, confirms the fact that the selected processes in these networks turn out to be age-dependent in a healthy population). It is also interesting that the *embryonic organ morphogenesis* process turned out to be significant for S-Control and M-Control Networks, but not for the AGE-Control. This indicates that this process is not characteristic of development in a healthy population at an age, but is different for DS compared to a healthy population of any age (which most likely indicates unstoppable embryonic development processes in the DS group). At the same time, it can be seen that both the S-Control and M-Control Networks contain unique processes (S-Control: *central nervous system neuron differentiation, appendage morphogenesis, muscle organ development, kidney epithelium development*; S-Control: *epithelial cell differentiation, forebrain development, camera-type eye development, neuron migration*), which indicates that the networks (in addition to the general factor - AGE-dependence) contain unique characteristic of disease properties.

**Table 3.** Gene Ontology results for Nodes of S/M/DS/AGE-Control 1-Dimensional comparison. The Table shows the results of Gene Ontology for loci selected during one-dimensional analysis. In this case, the high significance of the processes is achieved only for the AGE-Control and for the M-control (which most likely indicates the distinguished differences only within the age difference) comparison. Thus, the parenclitic approach is more informative in GO terms

### Generalized parenclitic network analysis

In this section, we describe the generalized parenclitic analysis algorithm. The primary aim of the algorithm is to identify deviations of an individual and their features from the Control group area on the plane of two arbitrary features. As a result, the found interrelations of features for an individual are presented in the form of an undirected graph, in which the features correspond to the vertices of the graph, and the presence of a connection between the features is an edge between the nodes (which exists if the individual falls outside the Сontrol area on the plane of two corresponding features, and does not exist if the individual belongs to the Сontrol area). The input data structure is a set of pairs for each *i*-th individual: (*X_i_,Y_i_*), where *X_i_* – the vector of features, *Y_i_* - is the class of the *i*-th individual: (-2, -1, +1, +2): where the absolute value equals to 1 and denotes individuals used in the construction of the classifier (or Train set) and 2 - used only in the estimation of deviations (or Test set); negative values correspond to the Controls group, and positive values correspond to the Cases group.

For example, in our design of the M-Control network, the group of mothers (DSM) had labels '-1' (because it is the Control group in this design), the group of children with DS (DS) was labeled '1' (because it is the Case group in this design). The group of healthy siblings (DSS), which does not participate in network training in this design, was labeled '2' (since this group also cannot be assigned to Cases and Controls, the use of the label in this case is not required).

#### Construction of a network

The proposed generalized parenclitic algorithm demonstrates the possibility of using any machine learning classifier as the core of the algorithm for selecting paired features and constructing the 'Control' area on their plane. To demonstrate this, we used two approaches in the implementation of the library: geometric (linear SVM) and probabilistic (PDF). As an improved version of the probabilistic approach, we propose a PDF-adaptive algorithm (which allows for automatic selection of the best cutoff threshold (for separation of ‘Control' area) on each plane of features pair, based only on the required separation quality).

Consider three of these options of the kernel of the parenclitic method:

- **Kernel: PDF reconstruction.** To assess the deviations of an individual from the control group, we propose using the distance between the said individual and the group. This distance should be calculated as an estimate of the empirical Probability Density Function (PDF). In this case, Kernel Density Estimation (KDE) with a Gaussian kernel is built on the plane for each pair of features, which are, in turn, based on the Control group samples. The algorithm can be described by the following sequence of 4 steps:
  1. Construction of an estimate of the empirical PDF using KDE on samples from the Control group. Let us denote this function as PDF(*x, y*).

**Note:** *at this step, samples of the Сontrol group marked with the Y label ‘ -1’ are used (that is, the set of Сontrols, which is assigned to the Train set)*

2. Calculation of the distance between each individual and the probability density, where the distance is calculated using the formula:

$$d\left( x,y \right)=\int_{PDF\left( u,v \right)>PDF(x,y)} PDF\left( u,v \right)dudv,$$

(1)

where (*x, y*) - values of two selected features of the individual. This formula characterizes the probability of a randomly selected individual to fall into an area with a higher probability density of the Сontrol group than that for the selected individual. Thus, the closer the value of PDF(x, y) to the maximum value of the probability density PDF(x∗, y∗), the smaller the corresponding distance d(x, y) from the individual to the distribution of Controls.

3. Selection of the significance threshold of the deviation according to the level of probability. The distance values $d\left( x,y \right)$ lie in the range [0, 1], since by definition $d\left( x,y \right)$ is the probability of some event. Thus, to determine the cutoff threshold we should select a value in the range from 0 to 1. Due to the lack of precise rules for determining the threshold, we use a set of thresholds in the range of acceptable values. Such values, for example, can be selected from the list **thr** ∈ {0*.*1*,* 0*.*2*,* 0*.*3*,* 0*.*4*,* 0*.*5*,* 0*.*6*,* 0*.*7*,* 0*.*8*,* 0*.*9}.
4. Determination of whether an individual deviates from the Control group by comparing the calculated distance with a selected threshold. If the distance from the individual to the probability density: ($d\left( x,y \right)$ > **thr**) is exceeded, this is interpreted as a significant deviation of the individual from the Control group (which means assigning an edge to the individual on the pair of selected features).

**Note:** *A decision on the best threshold can be made based on Сontrol and Сase goups in the Train set by examining the quality of their characteristics in terms of class-separability.*

- **Kernel: Linear SVM classification.** The standard PDF algorithm has a long computation time, which limits its use for large datasets. A less resource-intensive approach is to use a classification method based on the support vector machine (SVM). In general, SVM can be used with different engines and options to get the best results (for example, radial SVM, which works similar to PDF in terms of detecting ‘cloudy’ areas detection of controls). In this implementation, we consider only the linear SVM approach.
- Due to the substantial time of computation required by the PDF kernel approach, we suggested a faster distance classifier method. The methylation data analysis of Down Syndrome blood shows that almost all interesting feature pairs represent simple point cloud configurations. The Support Vector Machine classifier is well applicable to such data. In the general Case, one can use different kernels inside the SVM and different parameters for the best results. To finalize all steps for the identification of deviations from the Control group one should:

1. Construct separating hyperplane by SVM algorithm on the samples with Y labels -1 and 1 (Controls and Cases from Train set).
2. Check if SVM gives a good separation of Cases and Controls. With a low classification accuracy of two classes, it is not possible to detect significant deviations from the Control group by this method, which makes it possible to skip such pairs of features. In addition, when classifying the quality of a cut, the number of edges is reduced, which reduces the complexity of further calculations.
3. Determine on which side of the hyperplane the individual is located (which means assigning an edge to the individual on the pair of selected features if it is located in the “Case” area).

**Note:** *The SVM classifier allows building the distance to the hyperplane, which can be translated into the distance from the Control area (and thus exactly satisfies the essence of the kernel of the parenclitic approach.). The main advantage of this approach is that the constructed separating hyperplane itself is the best separator for two groups and does not require additional selection of thresholds to identify the best boundary.*

- **Kernel: PDF-adaptive (the best threshold method).** The calculation of a distance in previous approaches helped us to reduce a two-dimensional problem to a one-dimensional one. These functions of distance allowed us to form a rule. The only essential complication in the PDF method is a choice of the threshold value. To solve this problem, this approach uses an automatic adaptive threshold selection algorithm. The choice of threshold determines the division of the plane into two sets, and an individual can be classified correctly or incorrectly into each of the said sets. Reducing the dimension from 2D to 1D by introducing a distance function allows us to reformulate the threshold selection problem as the problem of choosing the best separator between two sets of points on a one-dimensional axis. A similar problem arises when the classification is performed by using a decision tree. At each step of the algorithm, in order to find the best separator, all possible separators are enumerated and a classification quality measure is estimated for each them. As a result of enumeration, the optimal separator is selected, which gives the highest value of the quality measure. Such a measure of quality can be, for example, accuracy, gini impurity, and information gain. In the implementation of the PDF-adaptive approach, we use the information gain measure, also known as the Kullback–Leibler divergence. Thus, the third step of the PDF algorithm, which selects the threshold, is replaced by an algorithm for choosing the optimal separator that maximizes the information gain. Consider a set of distances *di*∈ [0,1] computed for the *i*-th individual and a target label *Y_i_* : (-1: Control from Train Set and 1: Case from Train set). Then the problem of finding **thr** can be stated as follows:

| max ⁡𝑰𝑮(𝑪𝑻([𝒅 > thr], ***Y***)), (thr ∈ [𝟎,𝟏]), | (2) |
| --- | --- |

where 𝑪𝑻(*prediction*, *Y*) calculates the contingency table for two vectors: *prediction* – for predicted labels, and *Y* – for target labels, 𝑰𝑮 is a function that calculates information gain for a given contingency table at a selected threshold **thr**.

**Note:** *The finite number of thresholds for which the contingency table differs allows us to reduce the problem of finding the maximum over all thresholds in the range to enumeration of a finite number of delimiters. Each separator can be defined as the average between two adjacent distances in ascending order: di<di+1.*

By default, in our paper, we use a PDF-adaptive kernel (due to the higher computational speed compared to the PDF-kernel and the ability to capture nonlinear structures in the data, unlike the linear SVM-kernel).

Also, the kernel can be selected based on the following considerations:

**Time complexity of algorithms:** this parameter affects the computation time. Regardless of the chosen kernel, the generalized parenclitic algorithm has the complexity O (n2 Kernel) (since the data with n features requires iteration over all their pairs), where one of the three considered kernels can act as the Kernel: SVM = *m*^2^*...m*^3^; PDF= $mL+m^{2}, (L{10}^{5})$; PDF-adaptive = *m*^2^ (where *m* is the number of individuals in the dataset, *n* is the number of features in the dataset);

**Data features:** linear SVM is applicable only for data in which the relationship between pairs of features can be determined using a linear separation of classes; PDF is well applicable for data of a complex spatial configuration of separable sets with the possibility of manual control of a single probability threshold for all pairs of features; PDF-adaptive is similarly applicable for data with a complex configuration in the feature space, while the choice of the threshold value is carried out automatically and individually for each pair of features;

**Kernel features:** linear SVM is a more stable kernel and it cannot distinguish classes consisting of several point clouds in the general case; in turn, the PDF kernel allows detecting separated groups within the same class and correctly calculate the distance between the individual and the distribution of the Control group of arbitrary complexity; PDF-adaptive acts similarly to the PDF kernel, except for the ability to select a threshold and reduce the robustness of the method against outliers, as well as the ability to detect finer relationships between features.

**Note:** *As mentioned in the main text of the manuscript, we emphasize that any machine learning classifier can be easily integrated into this implementation (this classifier can automatically decide whether an individual belongs to any class or, if the algorithm returns the probability of an individual belonging to a Case class, then this probability can be used because the distance and thresholds on it can be investigated separately)*

#### Additional information about kernels and parenclitic procedure

##### PDF

The algorithm of this kernel supposes that the input data consists of *d* = 2 features and *m* samples.

- First step is a calculation of the two-dimensional KDE function. It is represented by a sum of Gaussian functions and can be found as *O*(*d*^3^ + *m* · *d*^2^). Hence, in the Case *d* = 2, complexity is *O*(*m*).
- Next, the distance from each individual to the distribution is found by calculating the integral (1). The Monte Carlo method is proposed as an algorithm for estimating the value of the integral. This method does not provide a high level of calculation accuracy, but allows calculating the integrals for all subjects at a time. Consider the Monte Carlo method for PDF integration over the whole plane. The algorithm samples the values from the distribution and averages them. Let (*X_j_,Y_j_*) be a random variable, j = (*1,L)* identically distributed over the reconstructed distribution using KDE. Next, pairs (*x _j_, y _j_*) of these distributions are sampled and the average is calculated:

$$\frac{1}{L}\sum_{j=1}^{L} PDF(x_{j}, y_{j}) L\to\int_{R^{2}} PDF\left( , v \right)ddv=1$$

(3)

This step can be completed in $O(L\cdot d^{2}\cdot m)$ time, where *L* denotes the number of pairs to be sampled. However, to calculate the integral (1), it is necessary to consider only the part of the space corresponding to the excess of the values of the integrable function PDF of some constant value $PDF\left( x,y \right)$. In this case, the following formula can be used to calculate the integral

$$\frac{1}{L}\sum_{j=1}^{L} PDF\left( x_{j}, y_{j} \right)\cdot[PDF\left( x_{j}, y_{j} \right)>PDF\left( x,y \right)] L\to\int_{R^{2}} PDF\left( , v \right)ddv$$

(4)

Using this formula, the computation time for $m$ individuals will be $O(L\cdot d^{2}\cdot m^{2})$. To optimize this formula, we use the already calculated values of $PDF\left( x_{j}, y_{j} \right)$. To do this, let us order the pairs ($x_{j}, y_{j}$) by the value $p_{j}$=$PDF\left( x_{j}, y_{j} \right)$ in non-increasing order and calculate the cumulative sums $s_{j}=\sum_{k=1}^{j} p_{j}$. In this case, the computation time will be $O(L+L\cdot logL+L\cdot d^{2}\cdot m)$. Values for *m* individuals can be obtained using a binary search algorithm for the values $p_{j}$ , which correspond to the values of the partial sums $s_{j}$. This allows us to calculate the final integral. In total, this algorithm has time complexity$O(L+L\cdot logL+L\cdot d^{2}\cdot m + m\cdot loglog m )$. As a result of the experiments, the optimal value was found to achieve a balance between the accuracy of the integral calculations and the calculation time: *L* = 10^4^ or *L* = 10^5^.

After the calculation of the distance values, the determination of the presence of a connection for each individual is linear time from *m*.

##### SVM

The third approach to computing the kernel based on the linear SVM method does not use additional algorithms. This means that its computational complexity is equivalent to calculating the SVM on 2 features with m examples, which depends on the internal parameters and implementation. The total complexity of the algorithm with the SVM kernel for enumeration of all pairs is estimated at *O*(*n*^2^ · *m*^2^).

##### PDF-adaptive

The most time-consuming part of the algorithm is calculating the distances from an individual to the distribution. However, to determine the presence of an individual deviation from the distribution, it is possible to exclude the integration step from the algorithm by replacing it with a less computationally intensive one. To achieve this, let us assume that the distance decreases monotonically as the value of PDF increases, and for one value $p$=$PDF\left( x, y \right)$ , there is a single value $d=d\left( x, y \right)$. This, in turn, makes it possible to replace the problem of finding the threshold for distances *d* with the equivalent problem of finding the threshold for *p*-values. Thus, using *p*-values in the adaptive algorithm at the separation step makes it is possible to reduce the algorithmic complexity while maintaining the correctness of the algorithm. As a result, the calculation process is reduced to two steps: calculating the *p-*values for *m* individuals, which is estimated at *O*(*m*^2^), and the time to determine the delimiter based on the information gain is *O*(*m* log *m* + *m*). The total time complexity of the adaptive PDF algorithm for iterating over all pairs is *O*(*n*^2^ *m*^2^).

##### Feature Selection Algorithm

Building a network using the proposed algorithms creates a significant redundancy in the number of found pairs due to the presence of features that are one-dimensional classifiers of classes. Such features, when paired with an arbitrary feature, also show the separation of groups with high accuracy. However, in this case, the need for the second feature may not be due to its weak predictive power. Since only one feature dominates in the pair, this demonstrates the absence of a relationship between the traits.

To avoid the detection of unrelated pairs, it is necessary to exclude those features from the list that give a high quality of classification in the one-dimensional case. Consider the problem of one-dimensional classification of two groups: for the *i*-th individual, the input features is *X_i_* and the target label is *Y_i_*. As a classifier, we use a simple threshold by the value of the attribute (if *x_i_*<**thr** then Control, otherwise the Case). Then, similarly to the separation algorithm in the PDF-adaptive method, we choose the threshold that gives the maximum value of information gain:

| max ⁡𝑰𝑮(𝑪𝑻([*x* > thr], ***Y***)), (thr ∈ [𝟎,𝟏]), | (5) |
| --- | --- |

As a result of the enumeration, the separator **thr** will be selected and the value of classification accuracy (**ACC**) will be determined for it. The *max_score_1d* parameter defines the maximum allowable accuracy value at which the feature will not be excluded from the list (**ACC** < *max_score_1d*, default *0.75*). Features with a higher classification accuracy of the two groups will be excluded from further consideration of the pairs.

##### Thresholds analysis

The PDF-adaptive approach assumes two a priori selected parameters: *minimal score* (90% by default) and *1-dimensional maximal score* (75% by default). The *minimal score* helps reducing computations and exclude feature pairs that do not show good classification of groups. The second parameter is excluding features that give good one-dimensional classification of groups. Here, we shed light on the robustness of our findings to the small changes of those parameters.

The decreased *minimal score* leads to capturing of more CpG pairs that grow exponentially (Fig. 15 in Supporting Information). Despite this, the overall picture does not change. The results of separation between classes by number of CpG site nodes with the non-zero-degree were provided for 3 thresholds: 0.85, 0.9, 0.95 in Fig. 16 in Supporting Information). This figure shows that the distribution of these characteristics (in groups and between groups) does not change depending on the *minimal score*. Finally, we analyzed the influence of *1-dimensional maximal score* on the results. The similar results were obtained in that case: the increasing of the parameter leads to the higher number of identified pairs associated with difference of DS from their healthy siblings (Fig. 17 in Supporting Information). However, the relative group location on the plane and relative order remained the same regardless of parameter changes (Fig. 18 in Supporting Information). In accordance with this this analysis, all network configuration findings about relationships between family members remain unchanged in a wide range of parameters. The only change is a detection of a different number of CpG sites from the common list of pairs for different parameters.

#### Visualization

A visual representation of the process of building networks in each design is presented in Fig. 5 (A (M/S/AGE/DS), B (M/S/AGE/DS)), and Fig. 5 (C (M/S/AGE/DS)) presents networks for members of Family 1. In each construction, the Parenclitic Network Algorithm established a multidimensional boundary in the selected set (own for each construction) of attribute pairs, through which the set of the Control group was very clearly separated from the Case group. Samples of the Control group are located on one side of such a multidimensional border (that is, they have almost no edges in their networks), and the group of Cases turned out to be on the other side of this multidimensional border, which is indexed by an edge in each pair of selected signs, which serves as an indication that they lie on the other side of Control group. Thus, the illustrations of the networks for the participants in the Control (see, Mother network in Fig. 5 (C (M)), Sibling network in Fig. 5 (C (S)), Sibling network in Fig. 5 (C (AGE)), DS network in Fig. 5 (C (DS))), and Case groups (see DS network in Fig. 5 (C (M)), DS network in Fig. 5 (C (S)), Mother network in Fig. 5 (C (AGE)), Mother or Sibling network in Fig. 5 (C (DS))) clearly demonstrate the global topological differences of the selected attribute systems. It is interesting to consider the participants in the Test group in the first three constructions (see, Sibling network in Fig. 5 (C (M)), Mother network in Fig. 5 (C (S)), DS network in Fig. 5 (C (AGE))). These networks are characterized by an intermediate state (visually, they are not as dense as the Cases, but also not as scattered as the Controls). In order to investigate such intermediate states and establish their relationship with age, we turn to the analysis of the topological characteristics of the received networks for each participant in the dataset in each network construction.

#### Topological analysis of a network

Revealing the internal mechanisms of investigated complex systems can be done by applying network analysis. Insight into the structure of networks gives us a possibility to measure integral system state observed for different subjects. In addition, centrality measures locate the most important or interesting nodes (features for initial system) in the network.

In the previous section, the algorithms produced a network for each subject, where nodes represented features, and links — deviation by a couple of features from its expected state. In this study, the following metrics are considered:

- Metrics that associate a number with each node (feature): Betweenness, Pagerank, Closeness, Eigenvector centrality, Degrees;
- Metrics that associate a number with each link: Distance;
- Metrics that characterize graph as whole: Efficiency, Robustness, Max component size, Number of nodes, Number of links;
- Other metrics: Component sizes;
- Statistics for vector metrics: Minimal value, Maximal value, Mean, Standard deviation, Number of zeros;

Full metric description includes:

- Degree centrality is defined as the number of links incident upon a node. The degree can be interpreted in terms of the immediate risk of a node to catch whatever is flowing through the network (such as a virus, or some information)
- Betweenness centrality is a measure of centrality in a graph based on shortest paths. For every pair of vertices in a connected graph, there exists at least one shortest path between the vertices such that either the number of edges that the path passes through (for unweighted graphs) or the sum of the weights of the edges (for weighted graphs) is minimized. The betweenness centrality for each vertex is the number of these shortest paths that pass through the vertex. *The betweenness centrality measures how often each graph node appears on the shortest path between two nodes in the graph. Since there can be several shortest paths between two graph nodes *s* and *t*, the centrality of node *u* is:

$$c_{u}=\sum_{s,t\neq u} \frac{n_{st}(u)}{N_{st}},$$

$n_{st}\left( u \right)$is the number of the shortest paths from *s* to *t* that pass through node $u$, and $N_{st}$is the total number of the shortest paths from *s* to *t*. If the graph is undirected, then the paths from *s* to *t* and from *t* to *s* count only as one path (divide the formula by two).

- PageRank algorithm outputs a probability distribution used to represent the likelihood of an event, in which a person by randomly clicking links will arrive at any particular page. PageRank of undirected graph is statistically close to its degree distribution. *The PageRank centrality type results from a random walk of the network. At each node in the graph, the next node is chosen uniformly from the set of successors of the current node (neighbors for the undirected Case). The centrality score is the average time spent at each node during the random walk.
- Closeness centrality of a node is a measure of centrality in a network calculated as the reciprocal of the sum of the lengths of the shortest paths between the node and all other nodes in the graph. Thus, the more central a node is, the closer it is to all other nodes. The closeness centrality type uses the inverse sum of the distance from a node to all other nodes in the graph. If not all the nodes are reachable, then the centrality of node *i* is:

$$c_{i}=\left( \frac{A_{i}}{n-1} \right)^{2}\frac{1}{C_{i}}$$

$A_{i}$ is the number of reachable nodes from node *i* (not counting *i*), *n* is the number of nodes in *G*, and $C_{i}$ is the sum of distances from node *i* to all reachable nodes. If no nodes are reachable from node *i*, then $c_{i}$ is zero.

- Eigenvector centrality is a measure of the influence of a node in a network. Relative scores are assigned to all nodes in the network based on the concept that connections to high-scoring nodes contribute more to the score of the node in question than equal connections to low-scoring nodes. A high eigenvector score means that a node is connected to many nodes which themselves have high scores. The eigenvector centrality type uses the eigenvector corresponding to the largest eigenvalue of the graph adjacency matrix. The scores are normalized such that the sum of all centrality scores is 1.
- Distance to dividing line from SVM; or a PDF distance is defined in PDF kernel description.
- Component is a subgraph in which any two vertices are connected to each other by paths, and which is connected to no additional vertices in the initial graph. There are several components. Size of the component is the number of vertices in this subgraph.
- Efficiency of a network is a measure of how effective the information exchanges in the system are. Average efficiency is defined as:

$$\frac{1}{n(n-1)}\sum_{s\neq t} \frac{1}{d_{st}},$$

where $n$ denotes number of nodes and $d_{st}$ denotes the length of the shortest path between a node *s* and another node *t*.

- Robustness or the ability to withstand failures and perturbations is a critical attribute of many complex systems including complex networks. Robustness is calculated as the number of steps in the process of removing nodes with high degrees until links are presented in graphs.
- Analysis of network complexity: Evaluation of network analysis complexity affecting the computation time provided in this paper can be done using the algorithmic complexity. The main pipeline includes analysis of pairs of features and performing some special kernel estimation for them. So, the complexity of a framework algorithm is *O*(*n*^2^ *T_kernel_*(*m*)), where *m* denotes the number of samples and *n* — the number of features. The computational time of the kernel depends on implementation details. Therefore, the following is a detailed description of the PDF kernel calculation.

### Parenclitic implementation

Parenclitic is an open-source python library. It can be distributed through the PyPI repository. Current version 0.1.6 provides functionality described in this paper.

Our package provides 3 main features:

- Build, save, and load parenclitic network.
- Choose or create a kernel to identify edges.
- Compute network metrics based on the python-igraph package.

Performance tests are done on the following setup: 256Gb RAM, x2 CPU Intel Xeon Gold 6238 CPU @ 2.10GHz, 22 cores per socket, 2 threads per core, 88 threads in total. Computation time of a CpG dataset with 87 samples and 114674 features is 147562 seconds ∼ 41 hours.
