## Supplementary material for "Age-related trajectories of DNA methylation network markers: a parenclitic network approach to a family-based cohort of patients with Down Syndrome": Table 1, Supporting Information

### 2.1 KEGG Parenclitic

| **Description** | **S-Nodes**  **comparison (p)** | **M-Nodes**  **comparison (p)** | **DS-Nodes**  **comparison (p)** | **AGE-Nodes**  **comparison (p)** |
| --- | --- | --- | --- | --- |
| *Neuroactive ligand-receptor interaction* |  | 3.7E-5 |  | 3.5E-7 |
| *Circadian entrainment* |  |  |  | 8.5E-7 |
| *Axon guidance* | 1.0E-3 | 5.5E-4 |  | 8.1E-6 |
| *Hippo signaling pathway* | 2.9E-6 | 1.6E-5 |  | 1.0E-5 |
| *Glutamatergic synapse* |  |  |  | 1.6E-5 |
| *Calcium signaling pathway* |  | 1.4E-4 |  | 2.0E-5 |
| *GABAergic synapse* |  |  |  | 3.2E-5 |
| *Rap1 signaling pathway* |  |  |  | 4.6E-5 |
| *Morphine addiction* |  |  |  | 1.1E-4 |
| *Wnt signaling pathway* | 3.3E-5 |  |  | 1.2E-4 |
| *Signaling pathways regulating pluripotency of*  *stem cells* | 4.6E-5 | 1.1E-4 |  | 1.7E-4 |
| *Type II diabetes mellitus* |  |  |  | 2.9E-4 |
| *Nicotine addiction* |  |  |  | 4.7E-4 |
| *Retrograde endocannabinoid signaling* |  |  |  | 5.8E-4 |
| *Oxytocin signaling pathway* |  |  |  | 6.1E-4 |
| *Adrenergic signaling in cardiomyocytes* |  |  |  | 6.5E-4 |
| *Pathways in cancer* | 1.3E-5 | 4.1E-5 |  | 7.2E-4 |
| *cAMP signaling pathway* |  | 1.4E-4 |  | 8.8E-4 |
| *Estrogen signaling pathway* |  |  |  | 1.0E-3 |
| *Cocaine addiction* |  |  |  | 1.2E-3 |
| *Dilated cardiomyopathy* |  |  |  | 1.6E-3 |
| *Insulin secretion* |  |  |  | 1.8E-3 |
| *Dopaminergic synapse* |  |  |  | 1.9E-3 |
| *Maturity onset diabetes of the young* |  |  |  | 2.1E-3 |
| *Gap junction* |  |  |  | 2.9E-3 |
| *PI3K-Akt signaling pathway* |  |  |  | 3.4E-3 |
| *Arrhythmogenic right ventricular cardiomyopa-*  *thy (ARVC)* |  |  |  | 4.4E-3 |
| *Glycosaminoglycan biosynthesis - heparan sul-*  *fate / heparin* |  |  |  | 4.9E-3 |
| *Salivary secretion* |  |  |  | 5.0E-3 |
| *Cholinergic synapse* |  |  |  | 5.0E-3 |
| *Basal cell carcinoma* | 7.6E-4 |  |  | 9.1E-3  (Ben>0.05) |
| *Melanogenesis* | 5.4E-4 |  |  | 1.3E-2  (Ben>0.05) |
| *Proteoglycans in cancerc* | 1.4E-4 |  |  | 4.7E-2  (Ben>0.05) |

**Table 1.** Kyoto Encyclopedia of Genes and Genomes (KEGG) results (p value<0.05 and Benjamini <0.05).
