## Supplementary material for "Age-related trajectories of DNA methylation network markers: a parenclitic network approach to a family-based cohort of patients with Down Syndrome": Table 2, Supporting Information

### 2.2 GO Parenclitic

| **Term ID** | **Description** | **S-Nodes**  **comparison (log10 p)** | **M-Nodes**  **comparison (log10 p)** | **DS-Nodes**  **comparison (log10 p)** | **AGE-Nodes**  **comparison (log10 p)** |
| --- | --- | --- | --- | --- | --- |
| **GO:0007389** | *pattern specification process* | -5.684 | -5.8894 |  | -7.9355 |
| **GO:0003002** | *regionalization* | -5.1203 | -5.382 |  | -8.644 |
| **GO:0045165** | *cell fate commitment* | -4.9626 | -4.8861 |  | -5.0339 |
| **GO:0048562** | *embryonic organ morphogenesis* | -4.7282 | -5.3354 |  |  |
| **GO:0021953** | *central nervous system neuron differen-*  *tiation* | -3.804 |  |  |  |
| **GO:0001501** | *skeletal system development* | -2.9274 | -3.1411 |  | -2.5157 |
| **GO:0035107** | *appendage morphogenesis* | -2.859 |  |  |  |
| **GO:0048736** | *appendage development* | -2.8043 | -3.8576 |  | -3.3526 |
| **GO:0007517** | *muscle organ development* | -2.3705 |  |  |  |
| **GO:0072073** | *kidney epithelium development* | -2.1971 |  |  |  |
| **GO:0030855** | *epithelial cell differentiation* |  | -3.5217 |  |  |
| **GO:0030900** | *forebrain development* |  | -3.4096 |  |  |
| **GO:0043010** | *camera-type eye development* |  | -3.3543 |  |  |
| **GO:0001764** | *neuron migration* |  | -2.2874 |  |  |
| **GO:0097485** | *neuron projection guidance* |  |  |  | -4.2434 |
| **GO:0001655** | *urogenital system development* |  |  |  | -4.0381 |
| **GO:0007156** | *homophilic cell adhesion via plasma*  *membrane adhesion molecules* |  |  |  | -3.8404 |
| **GO:0030509** | *BMP signaling pathway* |  |  |  | -3.8404 |
| **GO:0072001** | *renal system development* |  |  |  | -3.8404 |
| **GO:0007423** | *sensory organ development* |  |  |  | -3.8404 |
| **GO:0061138** | *morphogenesis of a branching epithe-*  *lium* |  |  |  | -3.7958 |
| **GO:0001763** | *morphogenesis of a branching structure* |  |  |  | -3.6196 |
| **GO:0048608** | *reproductive structure development* |  |  |  | -3.4037 |
| **GO:0048705** | *skeletal system morphogenesis* |  |  |  | -3.3695 |
| **GO:0060173** | *limb development* |  |  |  | -3.3526 |
| **GO:0061458** | *reproductive system development* |  |  |  | -3.2725 |
| **GO:0030534** | *adult behavior* |  |  |  | -3.0895 |
| **GO:0043583** | *ear development* |  |  |  | -2.8072 |
| **GO:0019933** | *cAMP-mediated signaling* |  |  |  | -2.6638 |
| **GO:0007187** | *G-protein coupled receptor signaling*  *pathway, couple to cyclic nucleotide sec- ond messenger* |  |  |  | -2.4709 |
| **GO:0060541** | *respiratory system development* |  |  |  | -2.3026 |
| **GO:0007416** | *synapse assembly* |  |  |  | -2.1716 |
| **GO:0035270** | *endocrine system development* |  |  |  | -2.1445 |
| **GO:0007188** | *adenylate cyclase-modulating G-*  *protein coupled receptor signaling pathway* |  |  |  | -2.1072 |
| **GO:0060485** | *mesenchyme development* |  |  |  | -2.1072 |
| **GO:0007600** | *sensory perception* |  |  |  | -2.0474 |

**Table 2.** Gene Ontology results for Nodes of S/M/DS/AGE-Control Networks. It can be seen here that several processes that significant for the S-Control and M-Control Networks at the same time (e.g. *pattern specification process, regionalization, cell fate commitment, skeletal system development, appendage development*) turn out to be significant also for the AGE-Control (which, on a par with the study of network characteristics, confirms the fact that the selected processes in these networks turn out to be age-dependent in a healthy population). It is also interesting that the *embryonic organ morphogenesis* process turned out to be significant for S-Control and M-Control Networks, but not for the AGE-Control. This indicates that this process is not characteristic of development in a healthy population at an age, but is different for DS from a healthy population of any age (which most likely indicates unstoppable embryonic development processes for the DS group). At the same time, it can be seen that both the S-Control and M-Control Networks contain unique processes (S-Control: *central nervous system neuron differentiation, appendage morphogenesis, muscle organ development, kidney epithelium development*; S-Control: *epithelial cell differentiation, forebrain development, camera-type eye development, neuron migration*), which indicates that the networks (in addition to the general factor - AGE-dependence) contain unique disease properties characteristic.
