## Supplementary material for "Age-related trajectories of DNA methylation network markers: a parenclitic network approach to a family-based cohort of patients with Down Syndrome": Table 3, Supporting Information

## 2.3 GO 1D

| **Term ID** | **Description** | **S-Nodes**  **comparison (log10 p)** | **DS-Nodes**  **comparison (log10 p)** | **M-Nodes**  **comparison (log10 p)** | **AGE-Nodes**  **comparison (log10 p)** |
| --- | --- | --- | --- | --- | --- |
| **GO:0009986** | *cell surface* | -2.1981 |  |  |  |
| **GO:0007156** | *homophilic cell adhesion via plasma*  *membrane adhesion molecules* |  | -3.2689 |  | -18.51 |
| **GO:0007517** | *muscle organ development* |  |  | -3.6526 |  |
| **GO:0042391** | *regulation of membrane potential* |  |  | -3.4557 | -16.7595 |
| **GO:0023061** | *signal release* |  |  | -3.1597 |  |
| **GO:0006936** | *muscle contraction* |  |  | -3.1411 |  |
| **GO:0006935** | *chemotaxis* |  |  | -3.0779 | -9.1439 |
| **GO:0060078** | *regulation of postsynaptic mem-*  *brane potential* |  |  | -2.6991 | -11.0487 |
| **GO:0001764** | *neuron migration* |  |  | -2.5528 |  |
| **GO:0007610** | *behavior* |  |  | -2.5489 | -13.3862 |
| **GO:0061061** | *muscle structure development* |  |  | -2.3287 | -8.1952 |
| **GO:0007389** | *pattern specification process* |  |  |  | -18.0237 |
| **GO:0021953** | *central nervous system neuron dif-*  *ferentiation* |  |  |  | -17.2668 |
| **GO:0045165** | *cell fate commitment* |  |  |  | -15.7033 |
| **GO:0003002** | *regionalization* |  |  |  | -15.2857 |
| **GO:0043269** | *regulation of ion transport* |  |  |  | -15.1618 |
| **GO:0007423** | *sensory organ development* |  |  |  | -14.248 |
| **GO:0007600** | *sensory perception* |  |  |  | -13.6615 |
| **GO:0001655** | *urogenital system development* |  |  |  | -12.0915 |
| **GO:0007187** | *G-protein coupled receptor signal-*  *ing pathway, coupled to cyclic nu- cleotide second messenger* |  |  |  | -11.9508 |
| **GO:0060173** | *limb development* |  |  |  | -11.7471 |
| **GO:0048736** | *appendage development* |  |  |  | -11.7471 |
| **GO:0072001** | *renal system development* |  |  |  | -11.2255 |
| **GO:0050808** | *synapse organization* |  |  |  | -10.9626 |
| **GO:0002009** | *morphogenesis of an epithelium* |  |  |  | -10.5361 |
| **GO:1905114** | *cell surface receptor signaling path-*  *way involved in cell-cell signaling* |  |  |  | -10.3925 |
| **GO:0007188** | *adenylate cyclase-modulating G-*  *protein coupled receptor signaling pathway* |  |  |  | -10.1445 |
| **GO:0010817** | *regulation of hormone levels* |  |  |  | -10.0453 |
| **GO:0035270** | *endocrine system development* |  |  |  | -9.7077 |
| **GO:0010469** | *regulation of receptor activity* |  |  |  | -9.5513 |
| **GO:0001501** | *skeletal system development* |  |  |  | -9.0395 |
| **GO:0019932** | *second-messenger-mediated signal-*  *ing* |  |  |  | -8.8761 |
| **GO:0048514** | *blood vessel morphogenesis* |  |  |  | -8.5376 |
| **GO:0048880** | *sensory system development* |  |  |  | -8.3726 |
| **GO:0007416** | *synapse assembly* |  |  |  | -8.2518 |
| **GO:0030198** | *extracellular matrix organization* |  |  |  | -8.2097 |
| **GO:0006813** | *potassium ion transport* |  |  |  | -8.1046 |
| **GO:0019933** | *cAMP-mediated signaling* |  |  |  | -8.0462 |
| **GO:0051962** | *positive regulation of nervous sys-*  *tem development* |  |  |  | -8.0273 |

| **GO:0043062** | *extracellular structure organization* |  |  |  | -7.6556 |
| --- | --- | --- | --- | --- | --- |
| **GO:0001763** | *morphogenesis of a branching struc-*  *ture* |  |  |  | -7.5171 |
| **GO:0048863** | *stem cell differentiation* |  |  |  | -7.4737 |
| **GO:0048638** | *regulation of developmental growth* |  |  |  | -7.1752 |
| **GO:0006836** | *neurotransmitter transport* |  |  |  | -6.6289 |
| **GO:0010721** | *negative regulation of cell develop-*  *ment* |  |  |  | -6.3382 |
| **GO:0009914** | *hormone transport* |  |  |  | -6.3307 |
| **GO:0048608** | *reproductive structure development* |  |  |  | -6.1267 |
| **GO:0055123** | *digestive system development* |  |  |  | -5.9706 |
| **GO:0061458** | *reproductive system development* |  |  |  | -5.9508 |
| **GO:0048565** | *digestive tract development* |  |  |  | -5.6882 |
| **GO:0050803** | *regulation of synapse structure or*  *activity* |  |  |  | -5.4498 |
| **GO:0035637** | *multicellular organismal signaling* |  |  |  | -5.4437 |
| **GO:0007626** | *locomotory behavior* |  |  |  | -5.3401 |
| **GO:0060541** | *respiratory system development* |  |  |  | -5.0315 |
| **GO:0043410** | *positive regulation of MAPK cas-*  *cade* |  |  |  | -4.9666 |
| **GO:0099504** | *synaptic vesicle cycle* |  |  |  | -4.8125 |
| **GO:0001505** | *regulation of neurotransmitter levels* |  |  |  | -4.7878 |
| **GO:0071773** | *cellular response to BMP stimulus* |  |  |  | -4.1701 |
| **GO:0015850** | *organic hydroxy compound trans-*  *port* |  |  |  | -3.5866 |
| **GO:0001101** | *response to acid chemical* |  |  |  | -3.3683 |
| **GO:0006874** | *cellular calcium ion homeostasis* |  |  |  | -3.1569 |
| **GO:0017156** | *calcium ion regulated exocytosis* |  |  |  | -2.7569 |
| **GO:0071229** | *cellular response to acid chemical* |  |  |  | -2.7424 |
| **GO:0097479** | *synaptic vesicle localization* |  |  |  | -2.6338 |
| **GO:0051588** | *regulation of neurotransmitter trans-*  *port* |  |  |  | -2.5354 |
| **GO:0032355** | *response to estradiol* |  |  |  | -2.508 |
| **GO:0061351** | *neural precursor cell proliferation* |  |  |  | -2.4376 |
| **GO:0090066** | *regulation of anatomical structure*  *size* |  |  |  | -2.2752 |
| **GO:0001503** | *ossification* |  |  |  | -2.2128 |
| **GO:0048511** | *rhythmic process* |  |  |  | -2.1032 |

**Table 3.** Gene Ontology results for Nodes of S/M/DS/AGE-Control 1-Dimensional comparison. Shown here are the results of Gene Ontology for loci selected during one-dimensional analysis. In this case, the high significance of the processes is achieved only for the AGE-Control and for the M-control (which most likely indicates the distinguished differences only within the difference of the age difference) comparison. Thus, the parenclitic approach is more informative in GO terms.
